# Ultrasound neuromodulation of an anti-inflammatory pathway at the spleen produces sustained improvement of experimental pulmonary hypertension

**DOI:** 10.1101/2023.03.28.534567

**Authors:** Stefanos Zafeiropoulos, Umair Ahmed, Alexandra Bekiaridou, Ibrahim T Mughrabi, Naveen Jayaprakash, Nafiseh Saleknezhad, Chrystal Chadwick, Anna Daytz, Izumi Sato, Yemil Atish-Fregoso, Kaitlin Carroll, Yousef Al-Abed, Marat Fudim, Christopher Puleo, George Giannakoulas, Mark Nicolls, Betty Diamond, Stavros Zanos

**Author notes:** **Corresponding author:** Stavros Zanos, MD, PhD 350 Community Drive, Manhasset, NY, 11030.

## Abstract

**Background:** Inflammation is pathogenically implicated in pulmonary arterial hypertension (PAH); however, it has not been adequately targeted therapeutically. We investigated whether neuromodulation of an anti-inflammatory neuroimmune pathway involving the splenic nerve using noninvasive, focused ultrasound stimulation of the spleen (sFUS) can improve experimental pulmonary hypertension (PH).

**Methods:** PH was induced in rats either by SU5416 (20 mg/kg SQ) injection, followed by 21 (or 35) days of hypoxia (SuHx model), or by monocrotaline (60 mg/kg IP) injection (MCT model). Animals were randomized to receive either daily, 12-min-long sessions of sFUS or sham stimulation, for 14 days. Catheterizations, echocardiography, indices of autonomic function, lung and heart histology and immunohistochemistry, spleen flow cytometry and lung single-cell-RNA sequencing were performed after treatment to assess the effects of sFUS.

**Results:** Splenic denervation right before induction of PH results in a more severe phenotype. In both SuHx and MCT models of PH, sFUS treatment reduces right ventricular (RV) systolic pressure by 25-30% compared to sham therapy, without affecting systemic pressure, and improves RV function and autonomic indices. sFUS reduces wall thickness, apoptosis, and proliferation in small pulmonary arterioles, suppresses CD3+ and CD68+ cell infiltration in lungs and RV fibrosis and hypertrophy and lowers brain natriuretic peptide. Beneficial effects persist for weeks after sFUS discontinuation and are more robust with early and longer treatment. Splenic denervation abolishes sFUS therapeutic benefits. sFUS partially normalizes CD68+ and CD8+ T-cells cell counts in the spleen and downregulates several inflammatory genes and pathways in nonclassical and classical monocytes, and macrophages in the lung. Differentially expressed genes in those cell types are significantly enriched for human PAH-associated genes.

**Conclusions:** sFUS causes dose-dependent, sustained improvement of hemodynamic, autonomic, laboratory and pathological manifestations in two models of experimental PH. Mechanistically, sFUS normalizes immune cell populations in the spleen and downregulates inflammatory genes and pathways in the lung, many of which are relevant in human disease.

## Introduction

Pulmonary arterial hypertension (PAH) is a relatively rare but fatal disease, characterized by abnormal contraction and proliferation of smooth muscle cells in pulmonary arterioles, which along with neointimal formation, lead to a progressive increase in pulmonary vascular resistance and development of right heart failure. Current PAH-specific treatments act primarily by promoting pulmonary vasodilation and, despite improving symptoms and functional capacity, they don’t alter the natural progression of the disease.^1,2^ Therefore, there is an urgent need for disease-modifying treatments. Chronic immune dysfunction and lung inflammation play a critical role in PAH pathogenesis by creating a microenvironment of exuberant cell growth and promoting intimal and medial hypertrophy.^3,4^ However, treatments targeting inflammation, including ligand traps and biologics, have only recently started being tested clinically.^5–7^ Thus, it remains unclear whether targeting inflammation could be an effective, possibly disease-modifying, therapeutic strategy in the treatment of PAH.

Inflammatory processes are regulated by local and systemic immune, hormonal and neural mechanisms, including autonomic neuroimmune pathways, which directly regulate immune cells through local release of neurotransmitters.^8,9^ A well-studied neuroimmune pathway involves preganglionic vagal and sympathetic nerve fibers synapsing at abdominal ganglia and postganglionic efferent, noradrenergic nerve fibers innervating the spleen.^9^ Upon activation of the pathway, norepinephrine is released from the splenic nerve in the splenic parenchyma, causing splenic T-cells to secrete acetylcholine, which acts on cholinergic receptors on splenic macrophages and suppresses release of pro-inflammatory cytokines.^9^ Modulation of this pathway with electrical vagus nerve stimulation (VNS) reduces inflammation in several disease models^10^, and, recently, in patients with rheumatoid arthritis.^11^ VNS was recently tested in a rat model of pulmonary hypertension (PH), improving pulmonary hemodynamics and suppressing cytokines and pathological markers of inflammation.^12^ However, the clinical significance of this approach may be limited, due to off-target effects and the upfront risk of surgical implantation of a medical device.^13,14^ An alternative approach for activating the neuroimmune pathway to suppress inflammation is focused ultrasound stimulation of the spleen (sFUS).^15^ sFUS is a non-invasive, organ-specific neuromodulation approach that suppresses acute^15^ and chronic systemic inflammation^16^, to a similar extent as VNS.^15^ This non-invasive approach may represent an anti-inflammatory treatment, adjunct to PAH-specific treatments, in the hemodynamically sensitive population of PAH patients.

## Methods

### A detailed description of materials and methods is provided in Supplementary material

Across several experimental cohorts, PH was induced in 98 male Sprague-Dawley rats (5-7 weeks, 150-200g, Charles River) using either the Sugen-Hypoxia-Normoxia (SuHxNx) model or the monocrotaline (MCT) model, as previously described (Table S1).^17,18^ Animals in each cohort were randomly assigned to receive either sFUS or sham treatment. Our sFUS delivery system consists of transducer, a matching network, an RF power amplifier and a function generator, described previously.^15,19^ sFUS (or sham) treatments were administered with animals anesthetized. All procedures were approved by the Institutional Animal Care and Use Committee of the Feinstein Institutes for Medical Research. During terminal procedures, right and left heart catheterizations were performed and tissues were collected for lung and heart histopathology, cytokines, and serum brain natriuretic peptide (BNP) measurements. Echocardiograms and ECG recordings were performed longitudinally. In a cohort of animals, splenic nerve denervation (SND) was performed 3 days before the induction of the SuHxNx model; in another cohort, SDN occurred right before initiation of ultrasound therapy. In single cell RNA sequencing of lung cells, known and canonical cell-type marker genes were used for the identification of cell types. Differentially expressed genes (DEGs) between treatment arms were determined for each cell type. Gene-set enrichment analysis was performed using hallmark pathways from the Molecular Signature Database to annotate DEGs for biological pathways as well as using human PAH-associated genes obtained from DisGeNET^20^ and the Comparative Toxicogenomics Database^21^ to establish human relevance. For flow cytometry, splenic cells were isolated by mechanical dissociation using a standardized protocol. Cells were labeled with fluorescently tagged antibodies and then were analyzed using BD LSR II (BD Bioscience) and FlowJo Software (FlowJo, LLC, Ashland, CA).

## Results

### Preemptive splenic denervation worsens PH phenotype

There is sparse clinical evidence that splenectomy is correlated with PAH.^22,23^ To investigate a possible correlation between splenectomy and PAH in humans, we performed a meta-analysis of clinical studies and found that splenectomy, encountered in 6% of all PAH patients, significantly increases the risk of developing PAH compared with other lung diseases (OR 11.11, 95% CI 1.34 – 92.08, Fig. S1). In experimental PH, splenectomy has been shown to worsen the disease phenotype but the underlying mechanism remains unclear.^24^ Because splenic innervation is involved in the regulation of inflammation, we tested whether part of the effect of splenectomy on PH phenotype may be mediated by that mechanism. Preemptive SDN, or sham surgery, was performed, and 3 days later animals were either injected with SU5416 and exposed to hypoxia (FiO2=10%) for 21 days and normoxia for 14 days (SuHxNx model; Fig. 1A) or maintained in normoxia for 35 days (Fig. 1B). Successful SDN was confirmed with tyrosine hydroxylase staining of the spleen (Fig. S2). Denervated animals with PH have increased right ventricular systolic pressure (RVSP) compared to sham-operated animals with PH (93.0 ± 15.7 mmHg vs. 70.2 ± 11.3, p=0.03, Fig. 1C), as well as increased RV end-diastolic pressure (RVEDP) (11.4 ± 0.9 mmHg vs 8.5 ± 0.3, p=0.03). In SDN animals with PH, wall thickness of small pulmonary arterioles (PAs) is increased (Fig. 1E), and RV function, assessed by tricuspid annular plane systolic excursion (TAPSE), is significantly impaired (Fig. 1F). Mean arterial pressure (MAP) is similar among all groups (Fig. 1D). In animals without PH, preemptive SDN has no effect on RVSP (Fig. 1C). Together, these results indicate that splenic innervation plays an important role in the development of experimental PH.

**Figure 1.**
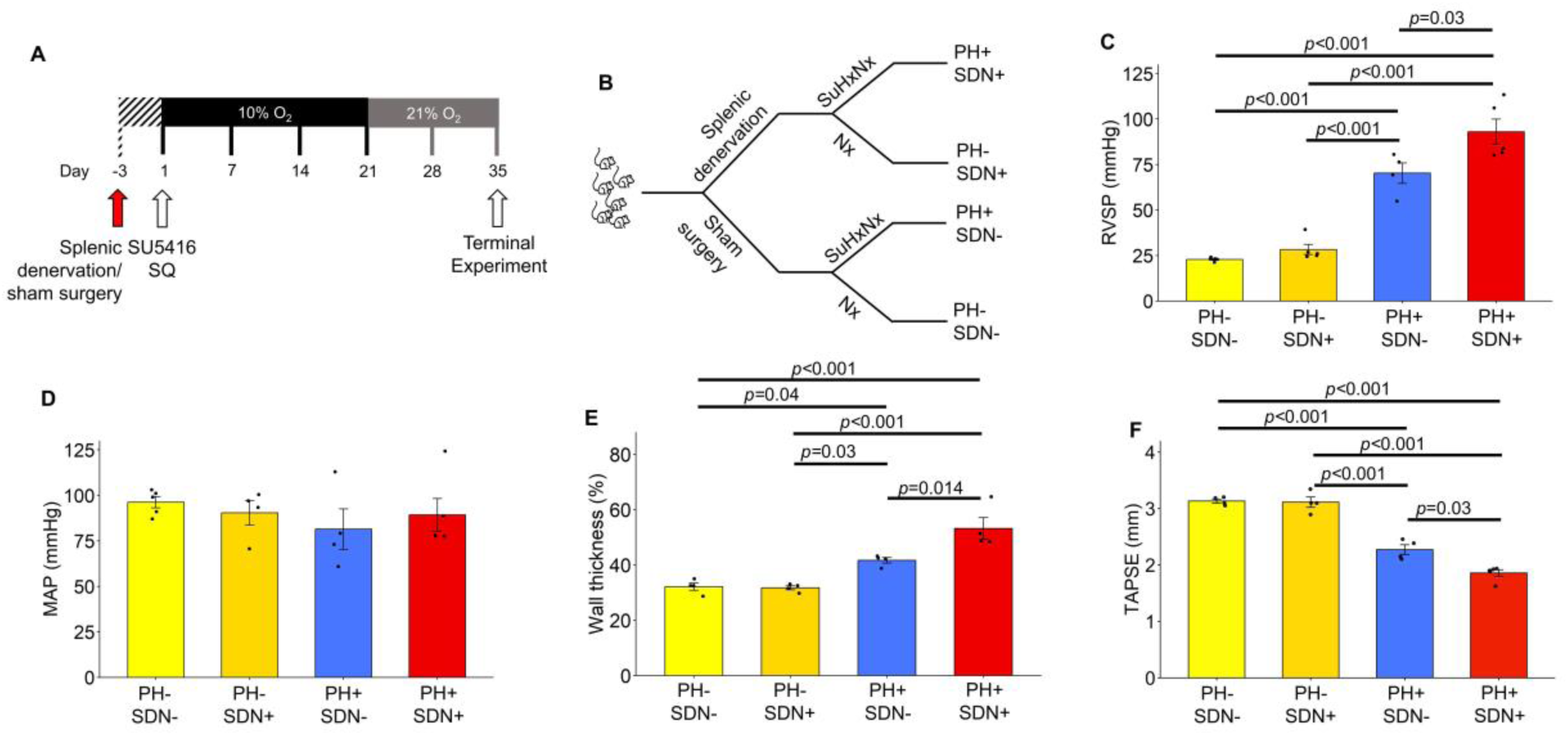
Pre-emptive splenic denervation worsens disease phenotype in a rat model of pulmonary hypertension. **(A)** Study design. On day −3, animals underwent splenic denervation (SDN), using a combination of surgical transection, local application of 100% ethanol and delivery of direct current to the splenic nerve, or sham-surgery. To induce pulmonary hypertension (PH), on day 1, animals were injected subcutaneously with Sugen 5416 (20 mg/kg) and, placed in a hypoxia chamber (10% FiO2) for 21 days, and re-exposed to normoxia for another 14 days; control animals remained in normoxia throughout the study period. A terminal experiment was performed on day 35. **(B)** The 4 experimental groups: animals with PH that received pre-emptive SDN (PH+/SDN+; n=5), control animals that received pre-emptive SDN (PH-/SDN+; (n=5), animals with PH that received sham surgery (PH+/SND-; n=4) and control animals that received sham surgery (PH-/SDN-; n=5). **(C)** Mean (±SE) right ventricular systolic pressure (RVSP) in PH-/SDN-, PH-/SDN+, PH+/SDN- and PH+/SDN+ animals; RVSP is significantly increased in PH+/SDN+ - compared to PH+/SDN-animals (p=0.03, one-way ANOVA with Bonferroni correction). **(D)** Mean (±SE) mean arterial pressure (MAP) in all 4 groups of animals; MAP is no different among all groups (p NS, one-way ANOVA with Bonferroni correction). **(E)** Mean (±SE) wall thickness in small (<50 μm in external diameter) pulmonary arterioles (PAs) in PH-/SDN-, PH-/SDN+, PH+/SDN- and PH+/SDN+ animals. Wall thickness was increased in PH+/SDN+ animals compared to PH+/SDN-animals (*p*=0.014, one-way ANOVA with Bonferroni correction). PA wall thickness was calculated by subtracting the internal diameter of the PA from its external diameter, divided by the external diameter. Ten PAs per animal were assessed. **(F)** Mean (±SE) tricuspid annular plane systolic excursion (TAPSE) in PH-/SDN-, PH-/SDN+, PH+/SDN- and PH+/SDN+ animals. Right ventricular function, as assessed by TAPSE, was worse in denervated animals compared to non-denervated animals with PH (*p*=0.03, one-way ANOVA with Bonferroni correction). Nx:Norrmoxia; SuHx:Sugen5416/Hypoxia;

### sFUS treatment improves pulmonary hemodynamics, reduces RV hypertrophy and restores indices of autonomic function

Because inflammation is implicated in PH^3^, and given our finding that splenic innervation in involved in PH pathogenesis, we hypothesized that sFUS treatment may suppress chronic inflammatory processes and ameliorate manifestations of PH. To test this hypothesis, rats with PH induced by SU5416 and hypoxia for 21 days and normoxia for 14 days (“Hypoxia-Normoxia cohort”, HxNx) received daily sessions of sFUS or sham, stimulation from day 22 to day 35. In addition, we tested sFUS in a more severe PH phenotype, by exposing animals to hypoxia for 35 days (“Hypoxia-Hypoxia cohort”, HxHx). In a subset of animals in the HxHx cohort, sFUS was delivered twice daily, for the same treatment period. A terminal experiment was performed on day 35 (Fig. 2A).

**Figure 2.**
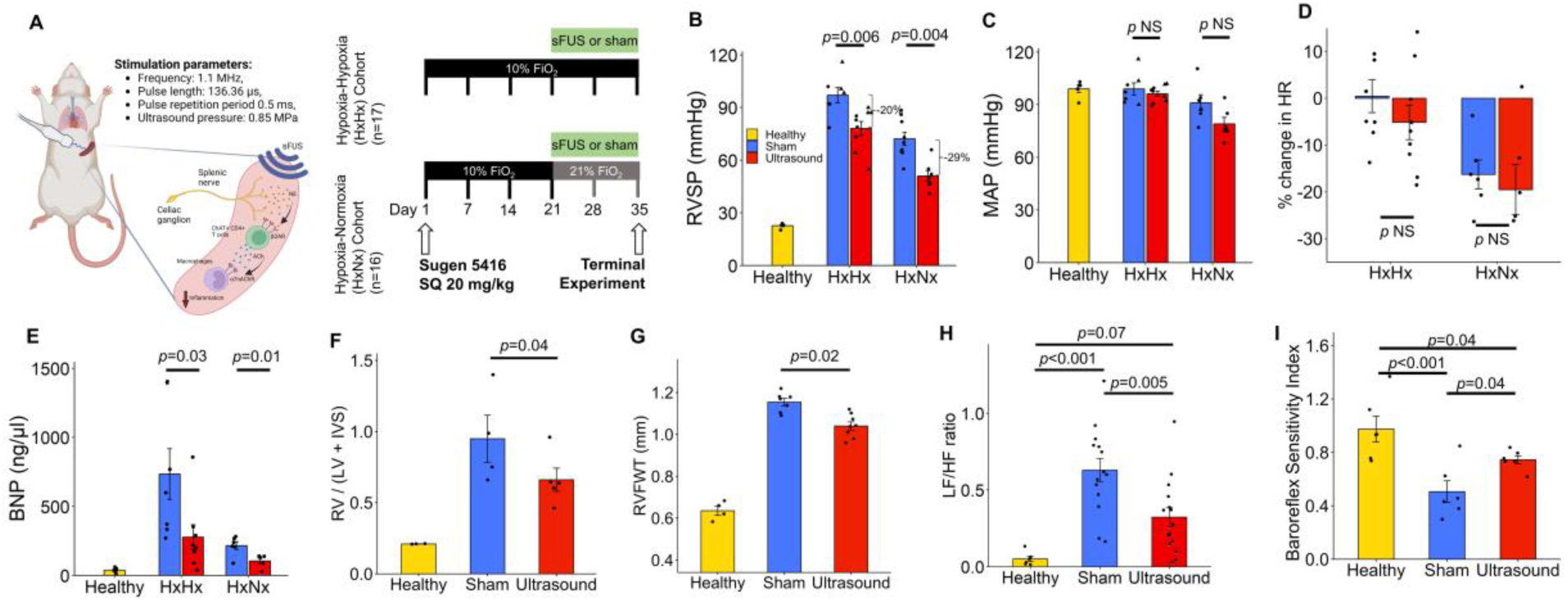
Focused ultrasound stimulation of the spleen improves pulmonary hemodynamics and reduces right ventricle hypertrophy in a rat model of pulmonary hypertension. **(A)** Study principle, design and timeline. Focused ultrasound stimulation of the spleen (sFUS) is delivered in Sprague-Dawley rats non-invasively at 0.83 MPa, pulsing frequency of 1.1 MHz and pulse repetition period of 0.5ms, parameters that have been shown to activate an acute anti-inflammatory response in rats.^15^ We used the SuHx model with either 21 days of hypoxia and then 14 days of normoxia (hypoxia-normoxia cohort, HxNx) or 35 days (hypoxia-hypoxia cohort, HxHx). Focused ultrasound or sham stimulation was delivered daily for 12 minutes under isoflurane anesthesia, starting at day 21, for 14 days. Right heart catheterization was performed at the end of treatment period. **(B)** Average (±SE) right ventricular systolic pressure (RVSP) in healthy, age-matched animals (n=5 animals; yellow bar), in the HxHx cohort: sham-treated animals (n=7), in sFUS-treated animals (n=9); in the HxNx cohort: in sham- and sFUS-treated (n=9, and n=7, respectively); RVSP is significantly reduced in sFUS-treated compared to sham-treated animals in both cohorts (p=0.006 and p=0.004, respectively, one-way ANOVA with Bonferroni correction). **(C)** Average (±SE) mean arterial pressure (MAP) in healthy animals, and in sham- and sFUS-treated animals, in the HxHx cohort and in the HxNx cohort; MAP is no different between sFUS- and sham-treated animals, in either cohort (p NS, one-way ANOVA with Bonferroni correction). **(D)** Mean (±SE) percentage change in heart rate (HR) between first day and last day of treatment, in the HxHx and HxNx cohorts (HxHx: −5.14% ±3.71 vs. 0.46±3.51; p =0.293, HxNx: −19.53% ±5.36 vs. −16.79±3.1, p=0.61, t-test). **(E)** Left panel: Mean (±SE) serum brain natriuretic peptide (BNP) on day 35 in HxHx cohort (p=0.03, t-test). Right Panel: Mean (±SE) BNP in HxNx cohort (p=0.01, t-test). **(F)** Mean (±SE) Fulton index, calculated as RV/(LV+IVS), in healthy (n=3), sham-treated (n=4), and sFUS-treated animals (n=5) (p=0.04, Kruskal Wallis test with Dunn’s Test for multiple comparisons). **(G)** Mean (±SE) Right ventricle free wall thickness (RVFWT) obtained via the parasternal short-axis axis at the mitral valve level in healthy (n=4), sham-treated (n=7), and sFUS-treated (n=8) groups (p=0.024, Kruskal Wallis test with Dunn’s Test for multiple comparisons). **(H)** The low-frequency to high-frequency (LF/HF) ratio is a metric of HRV: small LF/HF ratio is associated with a “healthier” sympathetic-parasympathetic balance. Mean (±SE) LF/HF ration in healthy (n=6), sham-treated (n=14), and sFUS-treated animals (n=15) on day 35 (sFUS vs sham: p=0.005, ANOVA test with Bonferroni correction). **(I)** Mean (±SE) baroreflex sensitivity index in healthy (n=6), sham-treated (n=6), and sFUS-treated animals (n=6) (ANOVA test with Bonferroni correction). BRS index was calculated as the absolute value of change in HR (|ΔHR|) over the change in systolic blood pressure (ΔSBP), before and after phenylephrine injection (25 ug/kg).

We found that sFUS treatment significantly reduces RVSP on day 35 by approximately 30%, compared to sham stimulation, in the HxNx cohort, and by about 20% in the HxHx cohort (Fig. 2B). Twice-a-day sFUS treatment has no different effect on RVSP than once-a-day treatment (Fig. 2B). Mean arterial pressure (Fig. 2C) and heart rate (Fig. 2D) are not affected by sFUS treatment. During sFUS sessions, no acute effects on pulmonary or systemic hemodynamics are seen in either healthy or hypertensive animals (Fig. S3), suggesting that the hemodynamic effect develops progressively over time. In both HxNx and HxHx cohorts, sFUS treatment results in lower right atrial pressure (RAP) (Fig. S4A) but does not affect RVEDP (Fig. S4B). Right ventricle dP/dt_max_ and RV dP/dt_min_, which reflect RV contractility and relaxation, respectively, and are themselves correlated with RVSP^25^, are reduced in sFUS-treated rats (Table S2), suggesting reduced RV workload.

Serum BNP levels are reduced in sFUS-treated animals, in both HxNx and HxHx cohorts (Fig. 2E). After returning to normoxia, sFUS-treated animals gain more weight than sham-treated animals compared to their baseline (32.45% ± 1.89 vs 22.13 ± 0.95; *p* = 0.002). Additionally, sFUS treatment results in reduced RV hypertrophy: Fulton index is significantly reduced in gross pathology (Fig. 2F), and RV free wall thickness is reduced in echocardiography (Fig. 2G). In these cohorts, sFUS treatment did not significantly improve pulmonary artery acceleration time (PAAT), PAAT/pulmonary artery ejection time (PAET) and TAPSE (Table S2, Fig. S5).

PAH is associated with autonomic imbalance, consisting of increased sympathetic and decreased parasympathetic tone.^26^ Commonly used markers of autonomic balance include heart rate variability (HRV) and baroreflex sensitivity (BRS)^27^, both of which are reduced in the PH animal model^28^ and in patients with PAH.^29^ Using spectral analysis of daily ECG recordings, we estimated the low-frequency (LF) and high-frequency (HF) components of HRV and calculated the LF/HF ratio, with lower ratios associated with improved sympatho-vagal balance (Fig. S6A). At the end of treatment, sFUS-treated animals have lower LF/HF ratio compared to sham-treated animals (Fig. 2H; Fig. S6B, independent comparison). In addition, sFUS-treated animals at end of treatment have a significantly smaller LF/HF ratio compared to beginning, whereas LF/HF ratio does not change in sham-treated animals (Fig. S6B, paired comparisons). Likewise, both LF and HF components improve in sFUS-treated but not in sham-treated animals (Fig. S6D-E, Table S3). The reduction in LF/HF ratio in sFUS-treated animals is first observed at day 31 and is sustained till the end of treatment (Fig. S6C). Across all animals in the HxNx and HxHx cohorts, the LF/HF ratio is significantly correlated with RVSP (Fig. S6F-H). To determine BRS, phenylephrine (PE) was injected intravenously and the ratio of change in heart rate over reflex change in systolic blood pressure was calculated (Fig. S7): the higher the ratio, the more sensitive the reflex. sFUS treatment for 14 days is associated with higher BRS index (Fig. 2I). Similar to the LF/HF ratio, BRS index is inversely correlated with RVSP (Fig. S6I).

*Taken together, these findings indicate that sFUS improves pulmonary hemodynamics without affecting systemic hemodynamics, reduces hypertrophy of the RV, and restores indices of autonomic function*.

### sFUS treatment improves vascular pathology and reduces inflammatory cell infiltration, apoptosis and proliferation in the lung

In the SuHxNx model of PH, lungs exhibit pathological abnormalities, including increased wall thickness in PAs.^17^ We performed morphometric analysis of PAs in H&E-stained lung samples (Fig. 3A) and found that sFUS-treated animals have reduced wall thickness in small PAs (Fig. 3B) but not in large PAs (Fig. 3C), compared to sham-treated animals. Using immunohistochemistry, we assessed lung infiltration from inflammatory cells, including monocytes and macrophages (CD68+) and T-cells (CD3+) (Fig. 3D and Fig. S8), and proliferation and apoptosis using Ki67 and TUNEL staining, respectively. In sFUS-treated animals, counts of both CD68+ and CD3+ cells are significantly reduced in both the lung parenchyma (Fig. 3E) and perivascular areas (Fig. 3F). Additionally, in sFUS-treated animals, Ki67-positive cells and TUNEL-positive cells (Fig. 3G-H) in the perivascular area are reduced compared with sham-treated animals (Fig. 3I-J).

**Figure 3.**
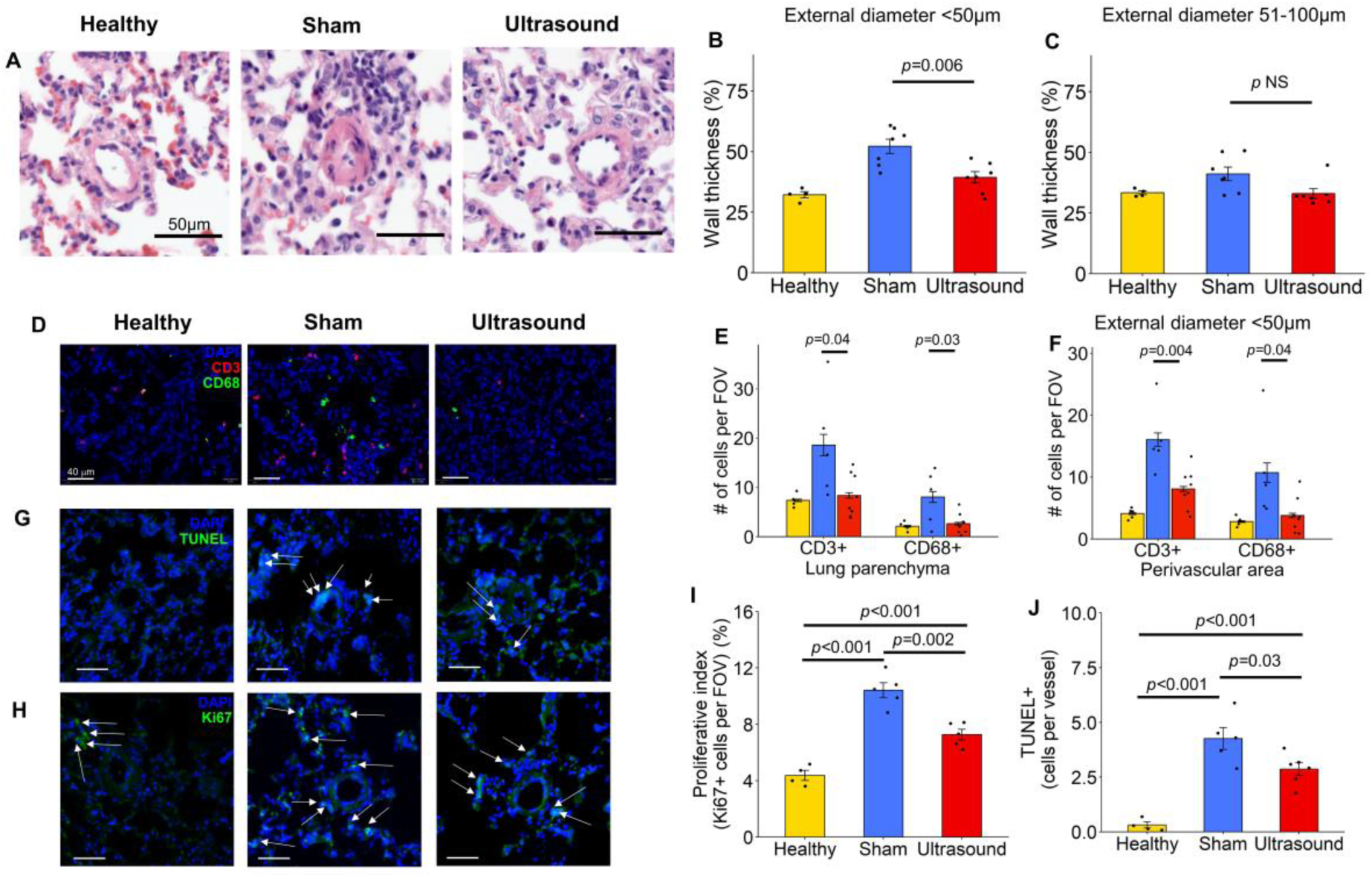
sFUS treatment improves vascular pathology and reduces inflammatory cell infiltration in the lung. **(A)** Examples of pulmonary arterioles (PAs) in healthy, sham-treated and sFUS-treated animals, shown in Hematoxylin and eosin (H&E) stain. **(B)** Mean (±SE) PA wall thickness in small PAs (<50 um in external diameter) in healthy (n=5 animals), sham-treated (n=7) and sFUS-treated animals (n=7) (p=0.006, one-way ANOVA test with Bonferroni correction; *p*=0.01, 2-way ANOVA with treatment group and animal ID as independent variables). PA wall thickness was calculated as in Fig. 1E. **(C)** Same as (B), but for PAs 50-100 um in diameter (p NS). **(D)** Images of lung sections stained with CD3+ antibodies (T-cells) and CD68+antibodies (macrophages), combined with nuclear DAPI stain in a healthy animal (left column), sham-treated animal (middle column), and sFUS-treated animal (right column). **(E)** Counts of CD3+ cells and CD68+ cells in healthy (5 animal), sham-treated (5 animals), and sFUS-treated animals (8 animals) (p values from one-way ANOVA tests with Bonferroni correction). Cells were counted in 10 random field of views (FOV) (150μm x 150μm) per sample. **(F)** Same as (E), but cells were counted in areas around <50 μm diameter PAs (perivascular area). (**G and H**) Images of lung sections stained with TUNEL and Ki67 antibodies antibodies **(I)** Proliferative index (Ki67+ cells) in small vessels and perivascular tissues in healthy (n=4), sham-stimulated (n=5) and sFUS treated rats (n=5) (p= 0.002 sFUS vs sham, one-way ANOVA with Bonferroni correction, 20 random FOV quantified per sample). **(J)** Number of apoptotic cells (TUNEL+ cells) per vessel in healthy (n=4), sham-(n=5) and sFUS-(n=6) treated rats (p=0.03 sFUS vs sham, one-way ANOVA with Bonferroni correction).

### sFUS improves outcomes in MCT-induced experimental PH

To test whether sFUS is effective in a second widely used model of PH, animals were injected with MCT (60 mg/kg, i.p.) on day 1 and received daily sessions of sFUS (or sham stimulation) from day 11 to day 24 (Fig. 4A). sFUS treatment significantly reduces RVSP by approximately 25%, compared with sham stimulation (Fig. 4B). Additionally, sFUS-treated animals have improved LF/HF ratio (Fig. 4C) and there is a trend towards reduced small PA wall thickness (Fig. 4D). However, PAAT, PAAT/PAET and TAPSE were similar in the 2 groups (Table S4). Overall, both the RVSP and the amount of pulmonary arteries occlusion in the MCT model were lower compared with the SuHxNx, as has been previously reported.^18^ The hemodynamic benefits of sFUS were replicated in the MCT model whereas the different experimental timeline and the less severe pulmonary vasculature remodeling might explain the less robust effect of sFUS on PA wall thickness.

**Figure 4.**
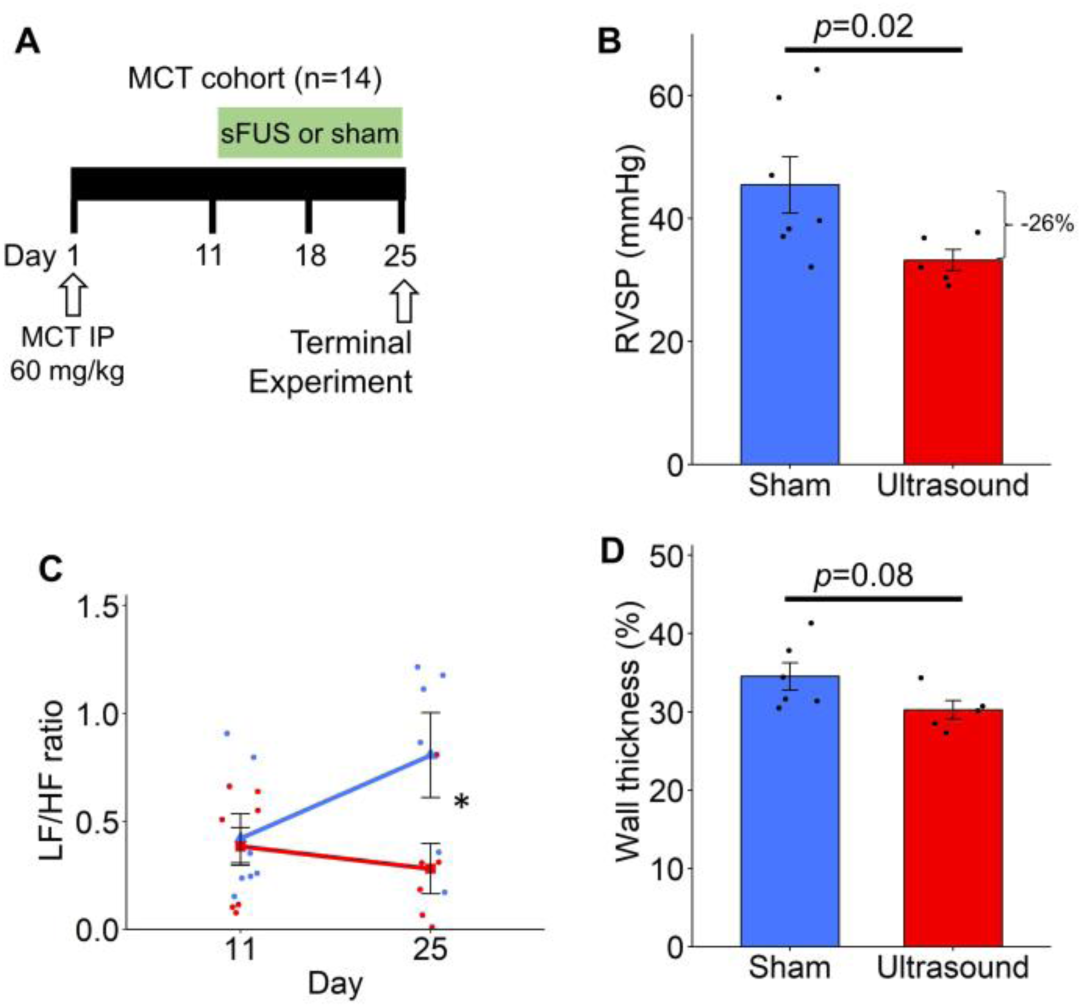
sFUS improves manifestations in the monocrotaline model of PH. **(A)** Experimental timeline in the “MCT cohort”. On day 1, rats (n=14) were injected with monocrotaline (MCT) (60 mg/kg) intraperitoneally and received daily sessions of sFUS or sham-stimulation from day 11 to day 24. The terminal experiment was performed on day 25. **(B)** Mean (±SEM) RVSP on day 25 (sham=7 vs. ultrasound=5; *p*=0.02, Wilcoxon rank sum test). (C) Mean (±SEM) LF/HF on day 11 and day 25 (sham=7, ultrasound=7, 2-way ANOVA on day 25 p=0.02). (D) Mean (±SEM) pulmonary vessel wall thickness percentage in the vessels <50 um in diameter in sham (n=6 animals) and ultrasound (n=5 animals) groups (t-test p=0.08).

### Therapeutic benefits of sFUS are sustained after discontinuation of treatment

To determine whether the therapeutic benefit of sFUS persists after discontinuation of treatment, in a separate cohort using the SuHxNx model, disease phenotype was assessed on day 49, 14 days after the end of treatment (“extended follow-up” cohort; Fig. 5A). In this cohort, sFUS-treated animals maintain most of the hemodynamic benefit: RVSP is still about 25% lower than in sham-treated animals (Fig. 5B). At that time point, RVSP values are higher than those on day 35 (i.e., as measured the HxNx cohort, Fig. 2B), reflecting the progression of disease in this model.^30^ RAP, LF/HF ratio and wall thickness of small PAs are reduced and RV function is improved in sFUS-treated compared to sham-treated animals (Fig. 5C-F).

**Figure 5.**
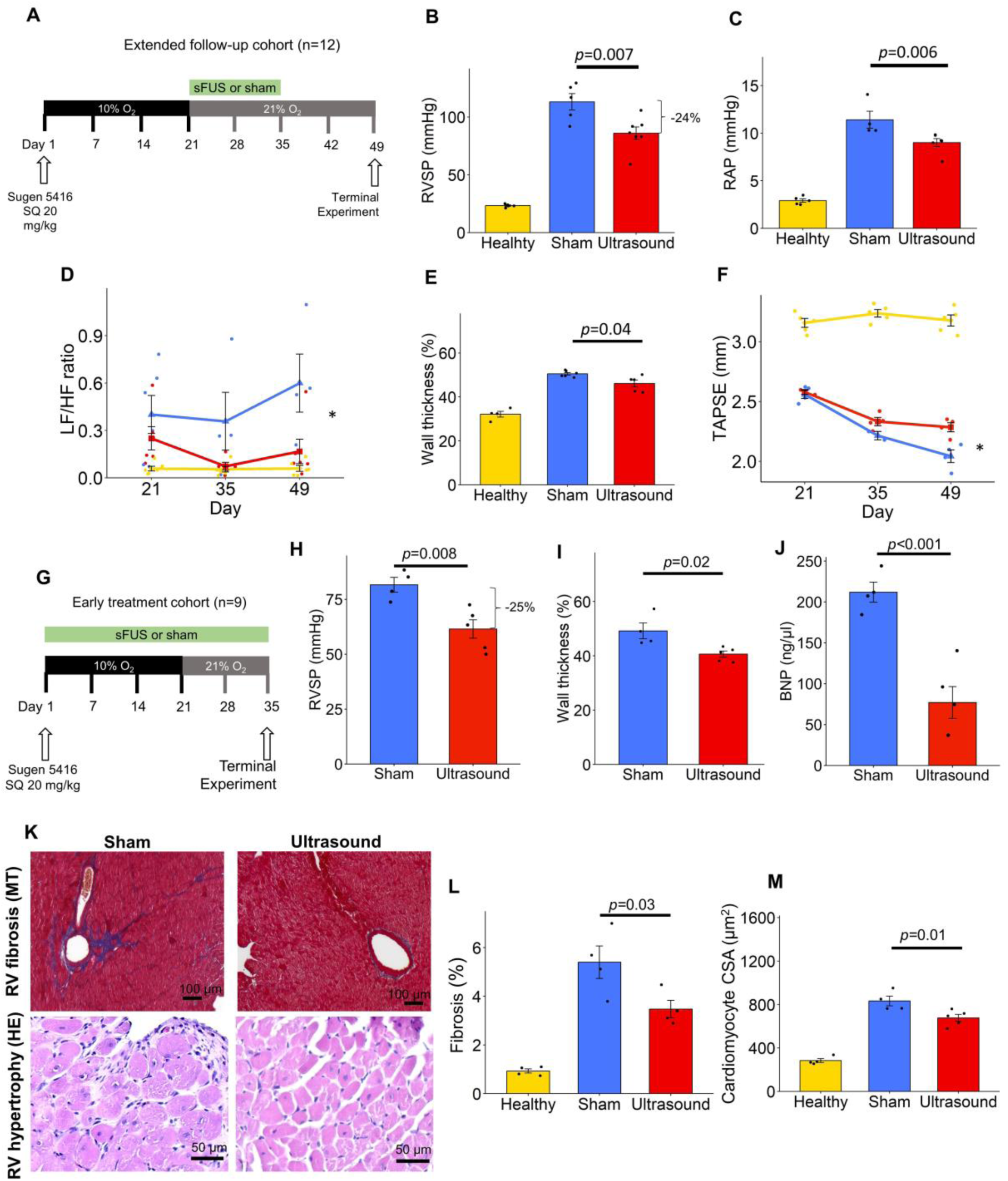
Sustained and dose-dependent therapeutic benefits of sFUS. **(A)** Experimental timeline in an “extended follow-up” cohort: PH was induced, and animals were treated with sFUS (or sham) for 2 weeks, similar to other cohorts; however, animals underwent terminal experiment 14 days after the end of treatment (day 49). **(B)** Mean (±SE) RVSP on day 49, 14 days after the end of treatment period (sham=5 animals vs sFUS=7; p<0.01, t-test). **(C)** Mean (±SE) right atrial pressure (RAP) at day 49 (sham=4 animals vs. sFUS=5; p=0.02, t-test). **(D)** Mean (±SE) LF/HF on days 21, 35 and 49 (sham=4 vs. sFUS=4). Asterisks at different time points represent significant difference between the 2 groups (*p*<0.05, repeated measures ANOVA). **(E)** Mean (±SE) wall thickness of small pulmonary arterioles (<50 um in diameter) assessed as in Fig 1E (sham=6 vs. sFUS=5; p=0.046, one-way ANOVA with Bonferroni correction). Using each vessel measurement, two-way ANOVA with treatment group and animal ID as independent variables, p<0.001 for treatment group, p=0.14 for animal ID). **(F)** Mean (±SE) TAPSE on days 21, 35 and 49 (sham=4, sFUS=4). Asterisks at different time points represent significant difference between the 2 groups (*p*<0.05, repeated measures ANOVA). **(G)** Experimental timeline in an “early and longer treatment” cohort: sFUS was initiated on the day SU5416 was injected; treatment was administered throughout the hypoxia and the normoxia period, for a total of 35 days, after which a terminal experiment occurred. **(H)** Mean (±SE) RVSP after 5 weeks of sFUS or sham treatment (sham=4 animals vs. sFUS=5 animals; *p*=0.008, t-test). **(I)** Mean (±SE) pulmonary vessel wall thickness percentage in small (<50 μm in diameter) pulmonary arterioles (*p*=0.001, t-test). **(J)** Mean (±SE) BNP levels (*p*<0.01, t-test). **(K)** Representative images of Masson trichrome (MT)- and hematoxylin and eosin (HE)-stained RV sections from a sham- and ultrasound-treated animal. **(L)** Mean (±SE) percentage area of fibrosis in RV (healthy=4, sham=4 vs. sFUS=4 animals; *p*=0.03, one-way ANOVA with Bonferroni correction). **(M)** Mean (±SE) cross-sectional area (CSA) of RV cardiomyocytes (healthy=4, sham=4 vs. sFUS=5 animals; p=0.01, one-way ANOVA with Bonferroni correction). CSA values were measured in at least 50 randomly chosen cardiomyocytes in each animal across the ventricle.

### Earlier, longer treatment with sFUS results in more robust functional and structural therapeutic benefits

To determine whether time of initiation and duration of treatment impact the therapeutic benefit, in a separate cohort using the SuHxNx model, sFUS (or sham) treatment started on day 1 and was delivered daily for a total of 35 days, including the 21 days of hypoxia (“early and longer treatment” cohort, Fig. 5G). In this cohort, sFUS robustly reduces RVSP, RAP and RVEDP (Fig. 5H, Fig. S9A-B). Autonomic indices, BNP levels, wall thickness of PAs are also improved (Fig. 5I-J, Fig. S9C-D) in sFUS-treated animals. sFUS improves RV hypertrophy and fibrosis (Fig 5K-M). IHC of the RV revealed less CD3+ cells by 75% and CD68+ cells by 30% in 2 sFUS-treated animals (Fig. S10) compared to 2 sham-treated animals. Additionally, TAPSE, free wall thickness, RV dP/dt_max_ and RV dP/dt_min_ (Fig. S9E-F, Table S5) are also improved.

### Splenic denervation eliminates the therapeutic benefit of sFUS

To determine whether the therapeutic effect of sFUS in PH is mediated by activation of the splenic nerve, we administered sFUS for 14 days in a separate cohort using the SuHxNx model that had received SDN, immediately before treatment initiation (day 21), (Fig. 6A-B and Fig. S11). In this “SDN cohort”, sFUS treatment does not result in lower RVSP compared to sham stimulation (Fig. 6C). No differences in heart rate (Fig. 6D), LF/HF ratio (Fig. 6E) or wall thickness of small PAs (Fig. 6F) are observed between sFUS- and sham-treated animals.

**Figure 6.**
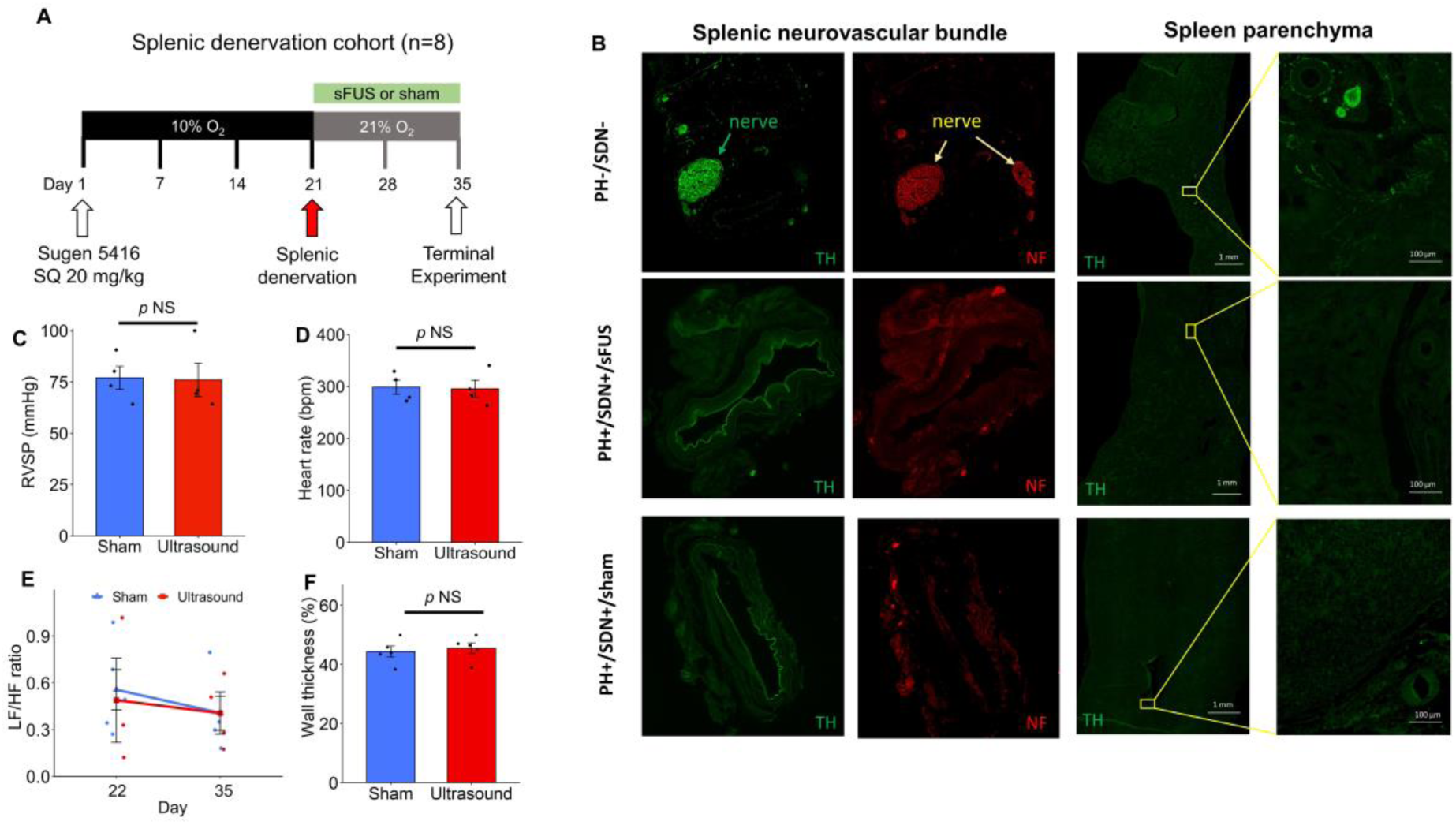
Splenic denervation eliminates the therapeutic benefit of sFUS. **(A)** Experimental timeline in a “splenic denervation” (SDN) cohort; at the end of hypoxia, animals undergo a survival procedure for SDN, followed by 14-days of sFUS (or sham) treatment and a terminal experiment. **(B)** Representative IHC images of adrenergic, tyrosine-hydroxylase (TH)-positive, neuronal fibers (stained together with neurofilament, NF, which stains all neuronal fibers) in the splenic neurovascular bundle (2 left-most columns) and in the spleen parenchyma (2 right-most columns) in a PH-/SDN-animal (top row), a PH+/SDN+/sFUS (middle row) and a PH+/SDN+/sham (bottom row). **(C)** Mean (±SE) RVSP of sham- and sFUS-treated denervated animals (Sham=4 vs. sFUS=4 animals; p nonsignificant, t-test). **(D)** Mean (±SE) heart rate (p nonsignificant, t-test). **(E)** Mean (±SE) LF/HF ratio (p nonsignificant, repeated measures ANOVA). **(F)** Mean (±SE) wall thickness (p nonsignificant, t-test).

### sFUS treatment partially restores CD8+ T and CD68+ cell populations in the spleen

Activation of the anti-inflammatory pathway at the spleen modulates the function of splenic T cells and macrophages, both essential steps in the suppression of systemic inflammation.^9^ To determine whether the anti-inflammatory mechanism of action of sFUS involves these cell populations, we performed flow cytometry in spleens of sFUS- and sham-treated animals with PH, as well as of healthy animals (Fig. S12). CD68+ cells were decreased in PH animals that received sham-stimulation compared to healthy animals (0.34% vs. 0.59%, p<0.001, Fig. 7A-B), and this was partially restored in the sFUS-treated animals (0.46% vs. 0.34%, p=0.02, Fig. 7A-B). Additionally, sham-treated animals with PH had higher number of CD3+CD8+ cells than healthy rats (14.16% % vs. 11.13%, p=0.004, Fig. 7C-D); sFUS partially restored this increase (12.10% vs. 14.16%, p=0.03, Fig. 7C-D), without affecting the total CD3+ and CD3+CD4+ cells (Fig. S13).

**Figure 7.**
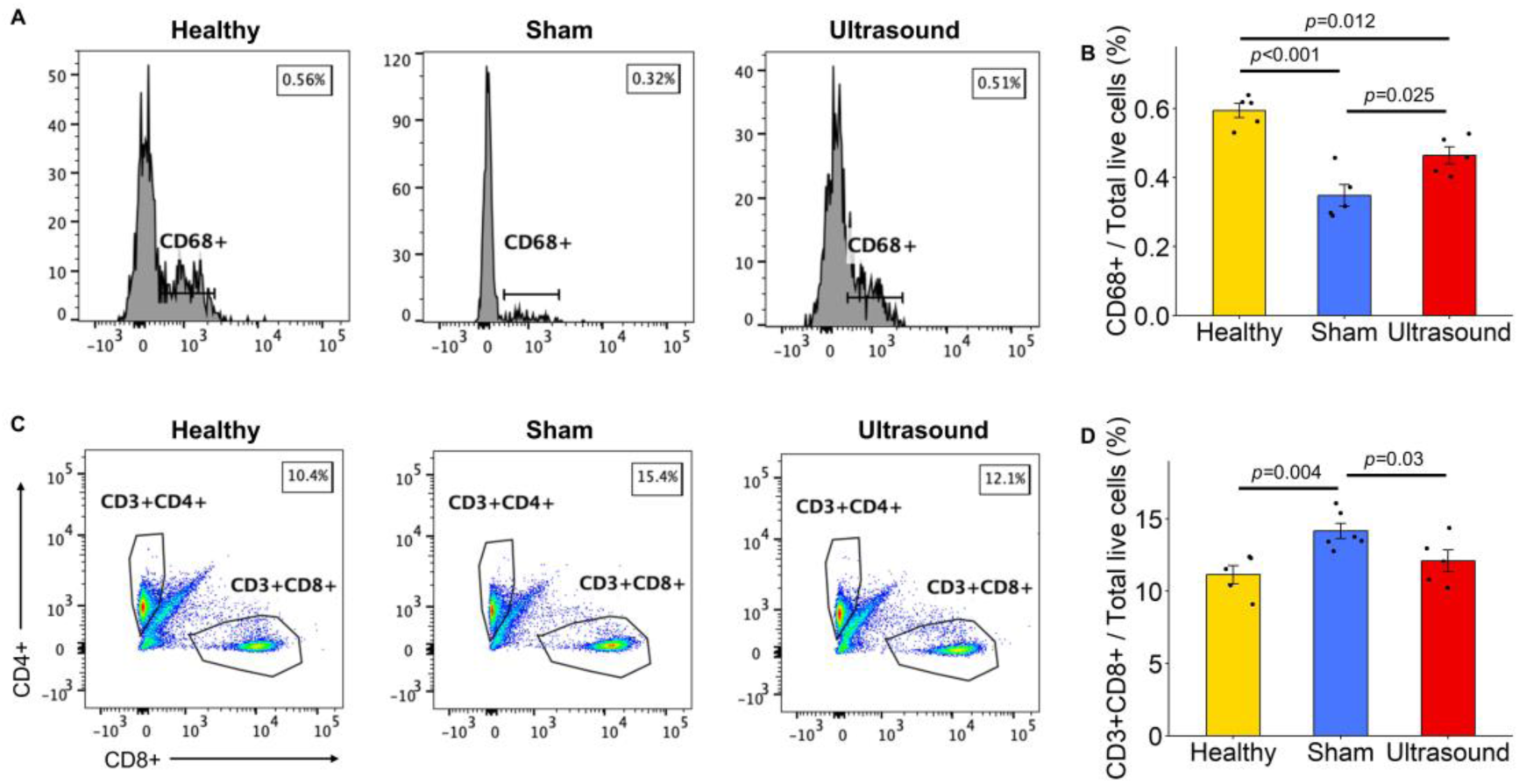
sFUS treatment partially restores CD8+ T-cell and CD68+ cell populations in the spleen. **(A)** Flow cytometry analysis of CD68+ cell population in the spleen. **(B)** Mean (±SE) percentage of CD68+ cells on day 35. Animals with PH (n=5) have decreased CD68+ macrophages compared to healthy animals (n=5) (one-way ANOVA with Bonferroni correction, p<0.001); sFUS-treated animals (n=5) have increased CD68+ cells compared to sham-treated animals (one-way ANOVA with Bonferroni correction, p=0.025). **(C)** Flow cytometry analysis of CD3+ T cell population and subsets. **(D)** Mean (±SE) percentage of CD3+CD8+ T cell population on day 35. sFUS-treated animals have decreased CD3+CD8+ cytotoxic T-cells compared to sham-treated animals (p=0.03, one-way ANOVA with Bonferroni correction).

In addition, we measured levels of circulating cytokines at the end of sFUS treatment. We found that sFUS reduces serum levels of interferon (IFN)-γ and interleukin (IL)-10 compared to sham-treatment; both cytokines are elevated in PH compared to healthy controls. In contrast, sFUS does not change levels of tumor necrosis factor (TNF) and IL-6, both of which are similar between PH animals and healthy controls (Fig. S14, Table S6). No differences were found between sFUS- and sham-treated animals in anti-inflammatory or pro-inflammatory cytokines in whole blood samples exposed to ex vivo lipopolysaccharide (23, 36) (Fig. S14, Table S7), lungs (Fig. S14, Table S9) and spleen homogenates (Fig. S14, Table S8), except of the increased anti-inflammatory IL-4 in sFUS-treated animals.

### sFUS treatment modulates transcription of inflammatory genes and pathways in immune cells in the lung

To screen for molecular mechanisms that could potentially mediate the therapeutic effects of sFUS in experimental PH, we performed scRNA-seq of lungs from 2 SuHxNx animals treated with sFUS and from 2 treated with sham stimulation. Sequencing profiled 7,870 cells after quality control steps, with even representation of groups. After clustering on the basis of transcriptomic similarity, 16 distinct cell types were identified, based on established markers for epithelial, stromal, lymphoid, and myeloid cell populations, and rare cell populations (Fig. 8A-B, Fig. S15A-B and S16). Lungs from the sFUS-treated animals have reduced proportion of both naive and CD8+ T-cells, and increased proportion of endothelial cells, compared with sham-treated animals (Fig. 8C, Fig. S15C). A total of 540 DEGs across 15 cell types were identified (false discovery rate <0.05) (Fig. 8D). We found that nonclassical monocytes (ncMono) and endothelial arterial 1 (EA1) cells have the most numerous DEGs (Fig. 8E) in the sFUS-treated animals. Pathway enrichment of DEGs revealed cell type–specific alterations of several inflammation-related pathways in different immune cell populations (Fig. 8F, Fig. S17). Notably, the most cell-specific downregulated pathways include the TNFa/NF-kB signaling in interstitial (iMF) and alveolar macrophages (aMF), ncMono, classical monocytes (cMono), T-regulatory cells, CD4+ T-naïve cells and endothelial capillary cells and the inflammatory response in iMF and cMono (Fig. 8G, Fig. S17). We further validated the downregulation of the NF-kB pathway in sFUS-treated animals by immunofluorescence, using nuclear staining for the NF-κB p65 in rat lungs of healthy, sham- and sFUS-treated rats (Fig. S18). We found strong nuclear p65 staining in the pulmonary arterial lesions of sham-stimulated rats with PH. Conversely, IFN-alpha response in ncMono and neutrophils are the most upregulated cell-specific pathways (Fig. 8H, Fig. S19). In a separate scRNA-seq experiment using animals from the early treatment cohort, the results were similar: inflammatory pathways in myeloid cells are downregulated in the sFUS-treated animals (Fig. S20). The same scRNA-seq analysis was performed for 2 healthy vs. 2 hypertensive rats: the TNF via NF-Kb signaling pathway in ncMonos is the most upregulated in animals with PH (Fig. S21), which is consistent with the published literature.^31^

**Figure 8.**
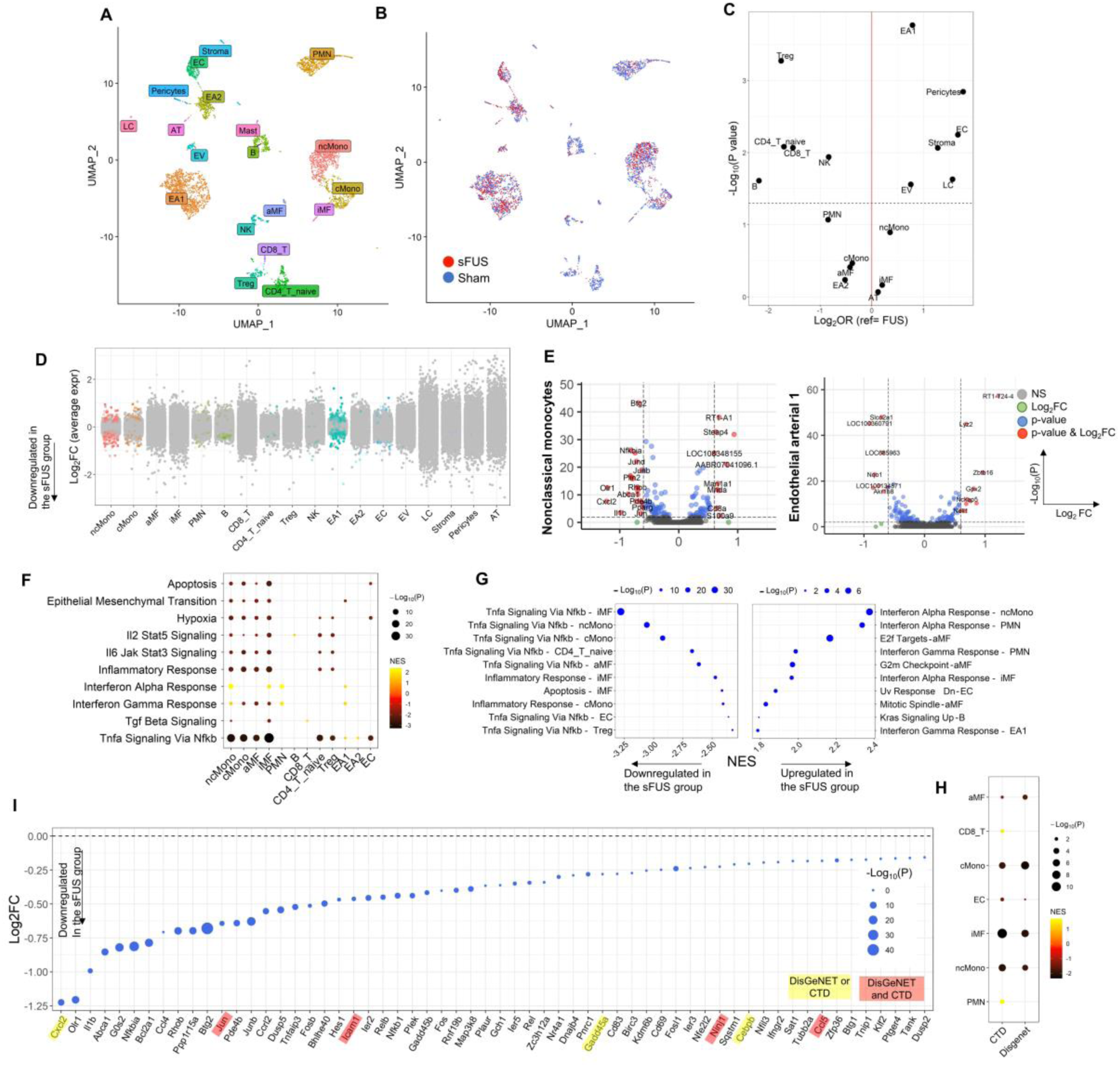
sFUS suppresses the expression of inflammatory genes and pathways in myeloid cells in the lung. **(A)** Uniform manifold approximation and projection (UMAP) plot showing lung cells from 4 rats with clusters labeled by cell type. **(B)** Uniform manifold approximation and projection (UMAP) plot showing lung cells from 4 rats with clusters labeled by stimulation status. **(C)** Plot showing comparisons of the clusters size between sFUS and sham-stimulated animals. X-axis represents the log2odds ratios (OR) of each cell type between sFUS and sham-stimulated animals while the y-axis the –log10(P-value). The dashed line represents a p-value of 0.05. **(D)** Jitter plot showing changes in gene expression for each cell type between sFUS and sham-stimulated rats. Each dot represents the differential expression MAST (Model-based Analysis of Single-Cell Transcriptomics) Log2-fold change of a gene. Dots indicating an adjusted p-value<0.05 are in color. The gray dots indicate values that were not significant (ns). **(E)** Volcano plots showing differentially expressed genes (DEGs) within the nonclassical monocytes (left) and endothelial arterial 1 (right) cells in the sFUS versus sham-stimulation groups in which the x-axis represents MAST (Model-based Analysis of Single-Cell Transcriptomics) log2 fold change and the y-axis indicates −log10(P-value). **(F)** Heatmap showing major cell type-specific pathway enrichment between sFUS and sham-stimulated rats using gene-set enrichment analysis (GSEA) (P<0.05) and 10 hallmark pathways from the Molecular Signatures Database that are known to be implicated in PAH on the y-axis. The dot size corresponds to −log10(P), and color represents the normalized enrichment score (NES) from GSEA, indicating upregulation (yellow) or downregulation (black). TNFa/NF-kB signaling was significantly downregulated across many cell types while interferon alpha response was the most upregulated pathway. **(G)** Dot plot showing the 10 most cell-specific downregulated (left) and upregulated pathways (right). **(H)** Heatmap showing most highly significant (p<0.05) enrichment of PAH genes of Disgenet and CTD databases in cell type–specific signatures using gene-set enrichment analysis, in which yellow indicates upregulation and black/dark brown indicates downregulation. The dot size represents −Log10(p-value). Significant upregulation of PAH genes from both databases was noted in myeloid cell types. **(I)** Dot plot showing MAST (Model-based Analysis of Single-Cell Transcriptomics) log_2_fold-change(Log2FC) of leading-edge genes accounting for the nonclassical monocytes downregulation of TNFa signaling via NfKb as determined by GSEA, in which the size and color tint of dots represent the −log_10_(P-value). Highlighted gene labels represent human pulmonary arterial hypertension–associated genes from either (yellow) or both (red) of the Comparative Toxicogenomics Database and DisGeNET Databases.

### Integrative analysis of rat scRNA-seq DEGs with human PAH-associated genes suggests possible clinical relevance of sFUS treatment

We performed an integrative analysis of rat scRNA-seq DEGs with human genes that are known to be associated with PAH. We identified genes implicated in human PAH from DisGeNET and the Comparative Toxicogenomics Database and tested rat DEGs for enrichment of these genes. We noticed significant enrichment of rat scRNA-seq DEGs in human PAH-associated genes from both databases, in myeloid cells, including iMF, cMono, ncMonos and aMF (Fig. 8H). We also noticed marked enrichment of human PAH-associated genes in the leading genes accounting for the downregulation of the TNFa via NF-kB pathway in the ncMonos (Fig. 8I).

## Discussion

In this study, we show for the first time that splenic innervation is implicated in the pathogenesis of experimental PH. Engagement of the splenic innervation using sFUS, a noninvasive neuromodulation method, leads to dose-dependent and sustained improvements in several clinically relevant hemodynamic parameters (RVSP, RAP), autonomic indices (HRV and BRS) and biomarkers with prognostic value (BNP). sFUS partially restores immune cell populations in the spleen and suppresses inflammatory cell infiltration in the lung, by altering expression of genes and pathways that are implicated in human PAH. To the best of our knowledge, this is the first report of preclinical efficacy of a noninvasive, ultrasound-based neuromodulation therapy in any cardiovascular or pulmonary disease, and more specifically in two models of PH, with indications for translational relevance to human PAH.

### Splenic innervation and sFUS in PH

Previous studies have suggested that splenectomy alters PH phenotype in animals^24,32^ and increases the risk of developing PAH in patients^22^, possibly because of increased platelet activation and chronic inflammation post-splenectomy.^33^ In our study, we find that animals that had received preemptive, selective SDN without splenectomy before the induction of PH have a worse phenotype, suggesting that altered splenic innervation may be another contributing factor and implicating splenic innervation in the development of PH. Electrical or ultrasound activation of the splenic innervation has been shown to suppress inflammation in animal models of acute endotoxemia, pneumonia, inflammatory arthritis, colitis and renal ischemia-reperfusion injury.^15,16,19,34,35^ Similar to previous studies^36^, we find that SDN blocks the anti-inflammatory effect of bioelectronic activation of the splenic innervation (Fig. 6). The splenic nerve is comprised of noradrenergic, efferent axons of ganglionic neurons in the celiac plexus, which regulate the function of immune cell populations in the spleen.^9^ Based on previous work^15^, it is likely that sFUS depolarizes axon terminals in the spleen to augment release of norepinephrine and activate the neuroimmune pathway—even though our study did not directly address this mechanism. Low intensity, focused ultrasound delivered to the brain produces mechanical waves that excite neurons via mechanosensitive ion channels, including TRPP1/2, TRPC1, and Piezo1.^37^ Similarly, FUS of the hepatoportal nerve plexus in animal models of diabetes restores glucose homeostasis via activation of TRPA1-positive, afferent nerve fibers.^38^ Notably, the abovementioned mechanosensitive ion channels are expressed in peripheral ganglia, in the vagus nerve and in some spinal nerves^39^ but the same channels are not expressed in other peripheral nerves, such as sciatic nerve, in which FUS does not elicit nerve responses.^40^ Furthermore, we find no evidence for an acute hemodynamic effect of sFUS in the systemic or the pulmonary circulation (Fig. S3), suggesting that the therapeutic effect of sFUS in PH is unlikely to be mediated by a direct effect on vascular tone or intravascular volume. Our finding that sFUS does not elicit acute hemodynamic effects has translational significance, as it suggests that this therapy may be safe in PAH patients, many of whom are easily compromised hemodynamically.

### Immunological effects of sFUS in PH

In a recent report, a single session of sFUS suppressed the rise in TNF after an acute inflammatory challenge, in the spleen and in whole blood of rodents.^15^ Similarly, 2 weeks of pretreatment with daily sFUS before an acute inflammatory challenge resulted in reduced, compared to sham, transcript levels of *Tnf* and associated genes^41^, suggesting that repeated sFUS may suppress gene expression across multiple molecular components of the innate immune response. Notably, it was recently reported that sFUS suppressed suppresses TNF release, in response to ex-vivo inflammatory challenge, from peripheral blood mononuclear cells in healthy human subjects^42^. Together with our finding that sFUS suppresses transcription of several inflammatory pathways in myeloid cells in the lung (Fig. 8), the evidence suggests that neuroimmune activation with sFUS may modulate widespread immune pathways, which in turn manifest as systemic and organ-specific anti-inflammatory effects, with potential application in PAH patients with chronic inflammation.

In our study, we find pathologic and transcriptomic evidence for reduced apoptosis, proliferation and inflammation in the lungs of sFUS-treated animals, including reduced counts of macrophages and T-cells. Macrophage and T-cell perivascular infiltrates and increased expression of inflammatory genes play a critical role in pulmonary vasculature remodeling in PAH^3,31^ and the effects on these cell populations could be partially responsible for the therapeutic benefit of sFUS. More specifically, in PH, blood-borne monocytes are deployed to the lung, where they differentiate into inflammatory interstitial macrophage.^43^ It is plausible that the deployment of splenic myeloid cells to the lungs plays an important role in PAH pathogenesis.^43^ Here, we find that rats with PH have increased number of macrophages (CD68+ cells) in the lungs and decreased CD68+ cells in the spleen, compared to healthy controls, and that sFUS treatment partially restores both these pathological alterations. This suggests that sFUS may promote retainment of macrophages in the spleen, partially explaining the reduced number in the lungs. Second, infiltration with CD8+ T cells can result in endothelial damage, disrupted apoptotic signaling and increased angiogenesis.^43,44^ We find that sFUS reduces CD8+ T cells in the lung, while reducing CD8+ T cells in the spleen. This suggests that CD8+ T cell differentiation and proliferation in PH may begin in the spleen^45^, where it is inhibited by sFUS.

Regulation of lung inflammation by the autonomic nervous system has received relatively limited attention in PAH. Augmentation of parasympathetic activity with electrical VNS or pyridostigmine results in improved hemodynamics, autonomic balance and attenuated cardiac and pulmonary vascular remodeling in experimental PH.^12,28^ In both cases, the beneficial effects of augmentation of parasympathetic activity were attributed mainly to reduced inflammation^12,28^, even though the mechanisms remain unclear. On the other hand, sFUS engages a well-defined, efferent neural pathway at the spleen, with subsequent anti-inflammatory effects, directly implicating autonomic control of inflammation in the therapeutic effects, as well as in the pathogenesis and progression of PH. Serum levels of the proinflammatory cytokine IFN-γ, a key regulator of macrophage activation through the JAK/STAT signaling pathway^46^, and IL-10 are reduced in treated animals, findings consistent with previously reported effects of sFUS on serum IFN-γ levels and IFN-γ gene expression in rats with inflammation and in patients with rheumatoid arthritis, respectively.^47^ We found no effect of sFUS on other proinflammatory cytokines in serum or in lung homogenates (Fig. S14), an expected result given that serum and tissue levels of almost all these cytokines were no different between healthy and PH animals (Fig. S14, Table S6-S9).

*Taken together, our findings that sFUS affects immune cells in the spleen and lung and has limited effects on circulating cytokines, suggest that sFUS regulates cell-mediated responses in PH*.

### Relevance of sFUS treatment in human PAH

We evaluated therapeutic effects of sFUS in two established, widely accepted animal models of PAH, using several clinically-relevant endpoints. The SuHxNx model in rats shares several common features with PAH in humans, including pulmonary hemodynamics^30,48^, pathological findings in the lung and heart^17^, and abnormalities in gene expression.^49^ In addition, efficacy of therapies tested with this model has been frequently associated with responsiveness in humans with PAH.^6,50^ In this model, sFUS reduces RVSP by 20-30% in animal cohorts with different severity of PH (Fig. 2), RAP, which is the strongest predictor of survival in PAH^49^, and serum BNP levels, a biomarker used in risk stratification of PAH.^51^

sFUS improves lung vascular pathology; it also improves heart pathology, including fibrosis, but it is unclear if that is a direct effect of sFUS on the RV, or a secondary effect of reduced afterload. Improved RV remodeling was evident in the early and longer treatment and the extended follow-up cohorts and, thus, might require longer period (i.e., more than the 14 days of treatment and follow-up as in the standard treatment cohort) of reduced afterload to become evident. Treatment with sFUS improves indices of autonomic tone that are commonly impaired in PAH patients^52^, similarly to other bioelectronic and pharmacologic therapies targeting the parasympathetic system.^12,28^ Our observation that both HRV and BRS correlate with pulmonary hemodynamics (Fig. S6) suggests that such noninvasive autonomic markers may prove useful in the evaluation and risk stratification of PAH patients. Unlike standard vasodilator therapies, the therapeutic benefits of sFUS persist for at least 14 days after treatment cessation, suggesting a potential disease-modifying effect of sFUS. Only one other study has demonstrated sustained therapeutic benefit after treatment cessation in experimental PH and, interestingly, that therapy targets activin-driven inflammation in pulmonary vascular remodeling.^6^ The development of disease-modifying therapies in PAH has been a challenge and sFUS warrants further investigation. Lastly, we observed that twice-a-day sFUS treatment does not further improve hemodynamics in PH, suggesting that once-a-day, or even a less frequent dosing scheme, may be sufficient.

We identified several inflammation-related pathways, including TNFa/NF-kB, IL-6/JAK/STAT, interferon gamma and inflammatory response, to be downregulated in sFUS-treated animals, with the TNFa/NF-kB pathway strongly downregulated in myeloid cells (Fig. 8). scRNA-seq studies of the lungs have shown that the same pathway is most prominently upregulated in both the SuHxNx and the MCT models, particularly in ncMonos and conventional dendritic cells, respectively.^31^ Additionally, TNFa signaling via NFkB is a major pathway associated with PAH in humans and DEGs in ncMonos in the SuHxNx model are most highly enriched for human PAH-associated genes.^31^ Interestingly, pharmacological inhibition of NF-kB improved pulmonary vascular remodeling, reduced immune infiltration in the lungs and preserved right ventricular function in SuHxNx rats, which are consistent with our overall findings.^53^ Notably, in our study DEGs in myeloid cells are significantly enriched for human PAH-associated genes; many of DEGs are also enriched in the leading genes of TNFa/NF-kB signaling pathway of ncMonos. We also observe upregulation of the IFN-alpha response by sFUS (Fig. 8), a pathway downregulated in the MCT model of experimental PH.^31^ Excess IFN signaling may contribute to PAH^54^ and exogenous IFN-alpha improved experimental PH and decreased proliferation in human pulmonary arterial endothelial and smooth muscle cells in vitro.^55^ These findings suggest that the anti-inflammatory and immunomodulatory effects of sFUS may have clinical relevance in human PAH and warrant further investigations.

### Limitations

Our study indicates that several inflammatory pathways are affected by sFUS treatment but provides no direct causal evidence that these effects are mediated through TNFa/NF-kB signaling or interferon-alpha. In addition, our study does not include detailed mechanistic investigations into how sFUS affects different functions of splenic immune cells, and how such alterations may impact PH pathogenesis. Therefore, additional preclinical and clinical studies are required to investigate the underlying mechanisms of sFUS treatment and further establish its real-world potential in PAH.

## Acknowledgments

The authors would like to thank Drs Valentin Pavlov, Lance Becker, Theodoros Zanos, Kevin Tracey, Sangeeta Chavan, Anne Davidson, Annette Lee, Christine Metz, and Peter Gregersen, and Mr. Houman Khalili for their critical appraisal of and comments on the manuscript.

