## Supplementary material for "Ultrasound neuromodulation of an anti-inflammatory pathway at the spleen produces sustained improvement of experimental pulmonary hypertension"

### Supplementary methods

#### Animal model of pulmonary hypertension and treatment cohorts

84 male Sprague-Dawley rats (5-7 weeks, 150-200g, Charles River) were injected subcutaneously with a vascular endothelial growth factor receptor inhibitor (SU5416; 20 mg/kg Tocris Bioscience and MedChem Express) dissolved in 0.5% carboxymethylcellulose, 0.9% sodium chloride, 0.4% polysorbate 80 and 0.9% benzyl alcohol, as described previously (1). The rats were placed in a hypoxic chamber with FiO2=10% (Biospherix Ltd) for 21 days (SuHx model); then they were re-exposed to normoxia. In the treatment cohort, 21 days after SU5416 injection, the rats were randomly assigned to either receive sFUS or sham treatment. Treatment period was 14 days and each daily session consisted of 12 minutes of sFUS (or sham stimulation). A subset of animals in the treatment cohort was not re-exposed to normoxia but remained inside the hypoxia chamber for a total of 35 days. In the early treatment cohort, rats underwent daily sFUS (or sham stimulation) during the whole study period, starting immediately upon SU5416 injection. On day 35 after SU5416 injection, animals in all but one cohort underwent a terminal experiment (Fig. 1A). In one cohort (extended follow-up cohort, Fig. 4A) animals were followed-up for an additional 14 days and the terminal experiment was performed on day 49.
12 male Sprague-Dawley rats (7-9 weeks, 250-350g, Charles River) were injected intraperitoneally with monocrotaline (60mg/kg, MedChem express) which was dissolved in 1 N hydrochloric acid and PH was adjusted to 7.4 with 1 N sodium hydroxide. Then, they received daily sessions of sFUS or sham-stimulation from day 11 to day 24. The terminal experiment was performed on day 25. All procedures were approved by Institutional Animal Care and Use Committee of the Feinstein Institutes for Medical Research. The sFUS system is a custom-made machine consisting of transducer, a matching network, an RF power amplifier and a function generator.

#### Focused ultrasound stimulation (FUS) of the spleen

The FUS system is a custom-made machine consisting of transducer, a matching network, an RF power amplifier and a function generator, as described previously (2). The transducer is acoustically coupled to the animal through a 6-cm-tall plastic cone filled with degassed water. The following stimulation parameters were used: pulsed sinusoidal waveform, 1.1 MHz pulse center frequency, 0.5 ms pulse repetition period, 200 mVpp amplitude, #Cycles:150, 0.83 MPa ultrasound pressure. A GE Vivid iq ultrasound system (GE Healthcare) equipped with a 12S probe (GE Healthcare) was used for the ultrasound scan before the first neuromodulation session. After the spleen was identified, the probe position was marked on the animal's skin and the FUS transducer was positioned on the marked area. During the daily 12-minute sFUS or sham-stimulation period, rats were anesthetized under 2% isoflurane. In the sham-stimulated animals, the probe was placed at the spleen, but the power remained off.

### Hemodynamic and physiological measurements

All terminal experiments were performed in the Center for Comparative Physiology (CCP) at FIMR. Animals were anesthetized at 5% isoflurane for 5 minutes in the induction chamber. Then, animals were placed in a supine position on a surgical platform and 2-3% isoflurane was delivered through a nose cone. 3-leads ECG were placed subcutaneously at the limbs of the animals to measure the heart rate. A rectal temperature probe was used to monitor body temperature. Body temperature was maintained at 37°C with a recirculating warming blanket. The depth of anesthesia was monitored by assessing heart rate, breathing rate and toe pinch. Under anesthesia, the neck region was shaved and sterilized. Invasive right heart catheterization was performed by making a midline incision in the neck, isolating the right exterior jugular vein, and then inserting and advancing the 2-F catheter (SPR-513, Millar) to the right ventricle (RV) through the vein. Invasive systemic arterial blood pressure was measured by insertion of a polyethylene-50 fluid filled catheter into the left carotid artery.

### Tissue harvesting

During the autopsy, lungs were intratracheally instilled with phosphate buffer solution. Lungs, hearts, and spleens were harvested. Lungs and hearts were fixed in 4% formalin, and the samples were kept at room temperature until histological and immunohistochemistry analysis. In some experiments, hearts were used to measure Fulton index as described below. Spleens were also harvested and stored in -80ºC freezer for cytokine analysis.

### Fulton Index

In some animals, the harvested hearts were used to measure Fulton Index [RV/ (LV + IVS)] to determine RV hypertrophy. The right ventricle (RV) was dissected from the left ventricle (LV) and interventricular septum (IVS), as described previously (3). The RV and LV+IVS weight was measured using a microbalance scale.

### Echocardiography measurements

Comprehensive echocardiography was performed at baseline, day 21 and day 35. Animals were anesthetized with isoflurane (2%), placed in supine position on a platform with a heating pad and subjected to echocardiography (GE Vivid iq equipped with a 12S-RS probe). During echocardiographic examination, we assessed pulmonary hemodynamics including estimation of mean pulmonary arterial pressure and pulmonary vascular resistance, pulmonary valve flow peak velocity, pulmonary acceleration time, pulmonary ejection time. We also assessed right heart structure and function, including right ventricular wall thickness, outflow tract diameter, and tricuspid annular plane systolic excursion. Measurements were performed off-line.

### Autonomic indices

Heart rate variability was evaluated with time- and spectral- domain analysis using the default rat algorithm of LabChart (ADinstruments). For the spectral-domain analysis, the low-frequency (0.2–0.75 Hz) and high-frequency (0.75–2.5 Hz) bands represented the sympathetic and parasympathetic components respectively and were expressed in both absolute (heart rate variability: $ms^2$), percent (%) and normalized units. The ratio between low-frequency and high-frequency components (low-frequency/high-frequency index) was considered a marker of the sympathovagal balance. Daily 12 min ECG recordings during the stimulation sessions of the treatment period were obtained. To evaluate the baroreflex sensitivity, we used phenylephrine (PE) (25 µg/kg) given by bolus injections (0.1 ml) to induce abrupt changes in systemic arterial pressure. A pressure catheter was placed in the left carotid artery to detect systemic arterial pressure. A second catheter was inserted in a right jugular vein to inject PE. Once the catheter was inserted, a small amount of saline was injected to confirm the catheter was functional. Injections were not repeated until the recorded parameters had returned to pre-injection levels. Baroreflex sensitivity index was calculated by identifying a 10 s window around the peak systolic blood pressure and corresponding heart rate after PE injection in a LabChart file (ADInstruments). Baseline values were calculated using a 10 s window before PE injection.

### Histology and Immunohistochemistry (IHC)

For histology and immunohistochemistry, left lungs were fixed in 4% buffered formalin, embedded in paraffin, and coronally sectioned at 5 µm thickness. For histology, the slides were stained with hematoxylin and eosin (H&E) for morphometric analysis of pulmonary arteries (PAs). Pulmonary arterioles were divided by outer vessel diameter into 2 categories: small (0–50 µm), large (51–100 µm). For each animal and each vessel category, 10 randomly selected vessels that were close to a round or oval shape were measured by an independent pathologist in a blinded fashion. The vascular wall thickness was calculated by subtracting the luminal (internal) diameter from the outer diameter circumference, as described previously. Wall thickness was expressed as a percentage, according to the following formula: 100*[(outer diameter) - (inner diameter)/outer diameter]. Completely obliterated vessels were also observed and included in the analyses. Hearts were also fixed 4% formalin, embedded in paraffin, sectioned cross-sectionally at 7 µm thickness and stained with H&E and Masson trichrome (MT). In the H&E staining, cardiomyocyte size for each ventricle was expressed as the average cross-sectional area of at least fifty transversally cut cardiomyocytes at the level of the nucleus, randomly distributed over the ventricles. In the MT, blue-stained tissue area was expressed as a percentage of the total surface area of RV. Images were collected using a BZ-X800 microscope (Keyence) at 20X magnification and a digital tablet camera. ImageJ software was used for measurements (4).

For multiplexed immunohistochemistry, the lungs were sectioned as described above. Slides were then cleared and processed for antigen retrieval as previously described (5). Protocols are also described in protocols.io (6-8). Briefly, slides were baked overnight and then taken though a series of xylene and ethanol to remove the paraffin. Slides were blocked with 4% BSA in PBS overnight at 4ºC, and then DAPI (4′,6-diamidino-2-phenylindole) stained (1 µg/ml), cover-slipped with anti-fade mounting media, and background imaged to collect DAPI, FITC, Cy3 and Cy5 filter channels. Slides were stained using a Leica Bond Autostainer with the following directly conjugated antibodies: CD68 (Abcam ab31630)-Cy3 and CD3 (Abcam ab16669)-Cy5. Imaging was conducted on a GE Healthcare IN Cell 2000 with Cell DIVE software (Leica Microsystems) for image registration, autofluorescence subtraction and correction. For the initial imaging step, a 10X objective was used, and the whole tissue imaged at 20x for DAPI and Cy3, followed by image stitching to create a composite image of the sample. The stitched image was then converted to a pseudo-H&E image as shown in Suppl. Fig S8 for visualization of histological features. QuPath v0.3.1 software was used for cell segmentation, classification and quantification (9). Between 1-3 lung sections were used to quantify cells for each animal. Immune cells were counted in two different sites, in the lung parenchyma and perivascular area. Ten regions in a slide were randomly selected using the same area measurement for both parenchyma and perivascular space across all the samples. For perivascular area, vessels with a diameter of <50 μm were used. Annotations from the regions selected had an optimal threshold set and mean fluorescent values were compared for the two different immune cell populations (CD3 and CD68). The number of proliferative cells was evaluated by anti-Ki67 antibody (1:250 dilution; ab16667, Abcam). Ki67 positive cells were counted around the pulmonary arteries (external diameter, <50 μm) in 20 random high-powered fields of view quantified per sample. Apoptotic cells were labeled using a TdT-mediated dUTP-biotin nick end labeling (TUNEL) in situ apoptosis detection kit (#25879, Cell Signaling Technology). TUNEL-positive cells around 10 random pulmonary arteries (external diameter, <50 μm) at ×400 magnification.

#### Cytokines and BNP analysis

Blood was collected in anti-coagulant heparin tubes during the terminal experiment and centrifuged at 2000 RCF for 10 min. Serum (supernatant) was collected and stored in a -80$^{o}$C freezer until further analysis. For tissue cytokines analysis, spleens were collected and stored in a -80 $^{o}$C freezer at the end of the terminal experiments. To homogenize the tissues, samples were taken out of the -80 $^{o}$C freezer, and 0.2 g of spleen were mixed with 1 ml of tissue reagent (T-PER Tissue Protein Extraction Reagent and Pierce Protease Inhibitor Tablets, Thermo Fisher Scientific). Tissues were then homogenized with Tissue Homogenizer (OMNI International), and centrifuged at 4700 RPM for 10 min. The supernatant was then collected and stored at a -80 $^{o}$C freezer until further analysis. Serum and tissue cytokines were analyzed using a V-PLEX Proinflammatory Panel 2 Rat Kit (K15059D-1, Meso Scale Diagnostics). Cytokines analyzed were TNF-a, IL-6, IFN-y, IL1b, IL-4. IL-5, IL-10, IL-13, and KC/GRO. Brain Natriuretic Peptide (BNP) was analyzed in the serum collected during the terminal experiment using ELISA kit (ab108815, Abcam). All steps for cytokines and BNP analysis were followed as recommended in the protocol by the manufacturer. All samples for cytokines and BNP analysis were performed in duplicates.

#### Splenic nerve denervation

In a subset of animals (n=8), we performed splenic nerve denervation (SND) on day 21. Rats were placed in a left lateral position; a subcostal incision was performed, and the spleen was exposed. After identification of the splenic neurovascular bundle, we isolated the splenic artery, the splenic vein, and the splenic nerve bundles, and then transected nerve bundles using a dissecting microscope. Then, we applied 100% ethanol locally on the splenic neurovascular bundle for 10 sec, and then we cuffed the splenic artery and delivered direct current of 4 to 5 V for 30 sec. At the terminal experiment, splenic arteries and spleens were harvested and stained with TH+ and confirmed successful SND (Suppl. Fig. S11). Using the same denervation protocol, we performed an additional cohort with pre-emptive SDN on day -3. Similarly, at the terminal experiment, spleens were harvested and stained with TH+ and confirmed successful SND (Suppl. Fig. S2).

#### Single-cell RNA sequencing

In a subset of animals (n=8), lungs were perfused, harvested, and were enzymatically dissociated into single cell suspensions, which was followed by scRNA-seq. Expression data was normalized, filtered, and clustered using the Seurat R package (R Foundation for Statistical Computing). Known and canonical cell-type marker genes were used for the identification of cell types (10). Cell-type proportions were quantified and compared between FUS- and sham-stimulated animals. Differentially expressed genes (DEGs) between treatment arms were determined for each cell type. Then, we performed gene-set enrichment analysis using hallmark pathways from the Molecular Signature Database to annotate DEGs for biological pathways as well as using human PAH-associated genes obtained from DisGeNET (11) and the Comparative Toxicogenomics Database (12) to establish human relevance.

### Flow cytometry

Single-cell suspensions were prepared by flushing the spleen with cold PBS then expressing the cells by manual compression. Red blood cells were lysed using 3 mL of ACK buffer for 6 minutes at room temperature. Cell counts and viability were determined using an automated count method (Nexcelom Cellometer Auto 2000 Cell Viability Counter Profiler). Splenocytes were suspended in flow cytometry buffer at 5x10 of cells and blocked with 1ug Fc block for 15 min. Cells were then stained with fluorophore-conjugated antibodies on ice for 30 min and washed twice with PBS containing 1% BSA and 0.02% sodium azide. The antibodies used were allophycocyanin (APC)-conjugated anti-rat CD68 (NB100-683APC), Brilliant Violet 605-conjugated CD3 (BD563949), Fluorescein isothiocyanate (FITC)-conjugated anti-rat CD4 (BD561833), Phycoerythrin (PE)-conjugated CD8 (BD559976). All labeled cell suspensions were fixed in 1% paraformaldehyde and stored protected from light at 4o C until analyzed by flow cytometry. The antibodies and the concentrations used in this study are presented in the Table. Cells were analyzed using BD LSR II (BD Bioscience) and FlowJo software (FlowJo, LLC, Ashland, CA).

| T-cells subsets panel | | Concentration |
|---|---|---|
| CD3 | BV-605 | 1:60 |
| CD4 | FITC | 1:60 |
| CD8 | PE | 1:60 |
| Macrophages subsets panel | | |
| CD68 | APC | 1:60 |
| CD86 | PE | 1:60 |
| CD163 | Alexa 405 | 1:60 |

Table. Flow cytometry antibodies

### Statistical analysis

All data expressed as mean ± standard error of the mean (SE). T-test or one-way ANOVA followed by post hoc test between healthy, sham- and sFUS-treated animals was used for normally distributed variables. Two-way ANOVA with treatment group and animal ID as independent variables was used for comparisons with multiple observations from each animal. Repeated measures ANOVA, followed by Bonferroni-corrected post hoc tests was used for longitudinal analysis of heart rate variability.

Mann-Whitney or Kruskal-Wallis test followed by the followed by the Dunn post hoc test between healthy, sham- and sFUS-treated animals was used for non-normally distributed variables. A p value $<0.05$ (after correction) was considered significant. All statistical analyses were performed with R Project for Statistical Computing (version 4.0.2).

### Supplementary figures

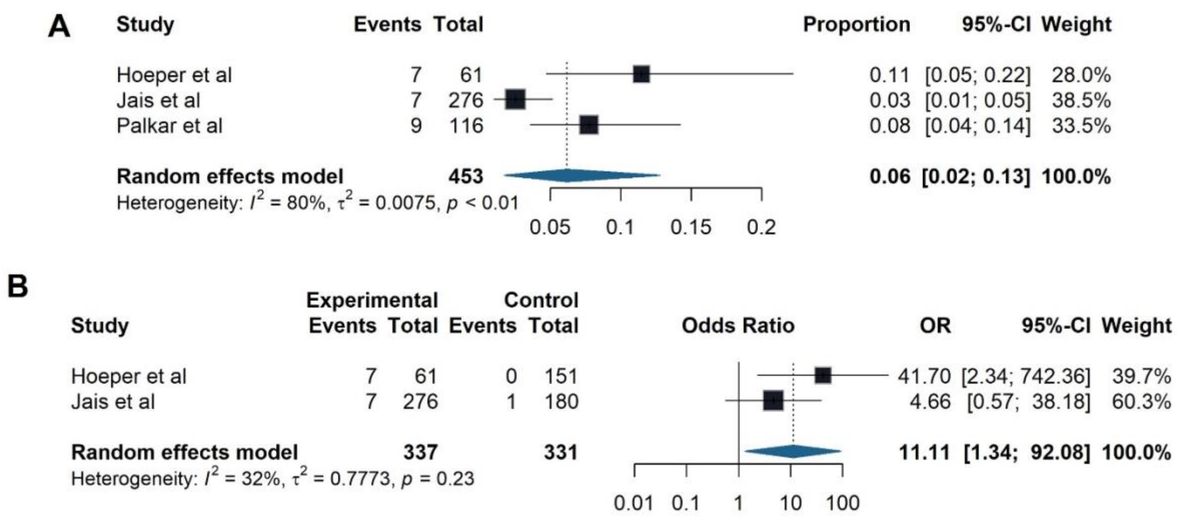


**Suppl. Fig. S1. (A)** Forest plot using a random effects single-arm model to calculate pooled proportion of splenectomy incidence among patients with pulmonary arterial hypertension. **(B)** Forest plot showing the odds ratio of splenectomy in developing pulmonary arterial hypertension compared to other lung diseases.

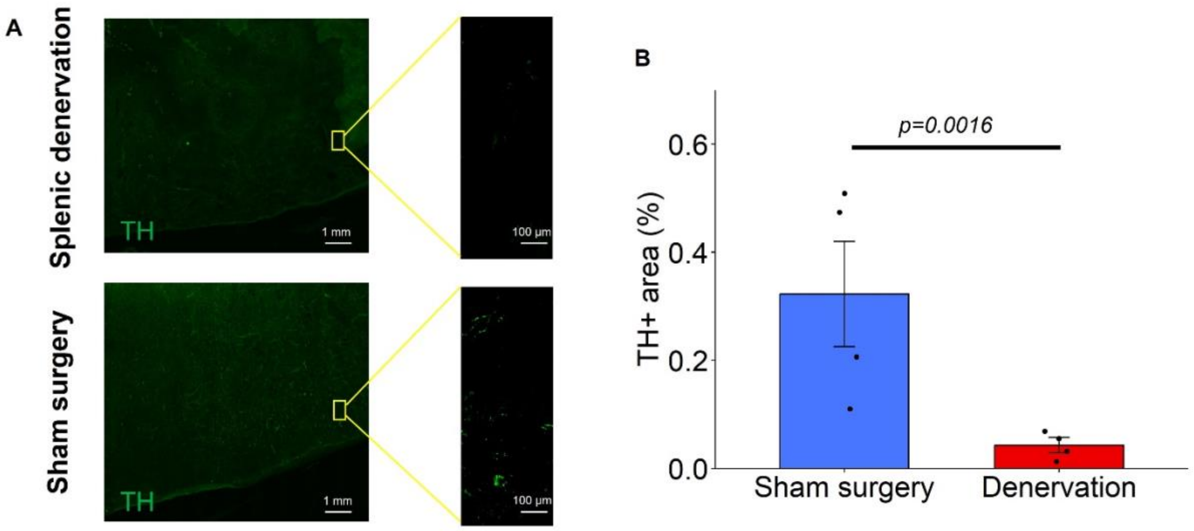


**Suppl. Fig. S2. (A)** Representative IHC images of adrenergic, tyrosine-hydroxylase (TH)-positive (left panels), neuronal fibers in the spleen parenchyma in an animal with denervated spleen (upper row) and an animal with sham-surgery (bottom row). **(B)** We quantified 5 same sized areas across the spleen, and we took the average of TH+ signal in animals that underwent sham-surgery (n=4) and denervated animals (n=4).

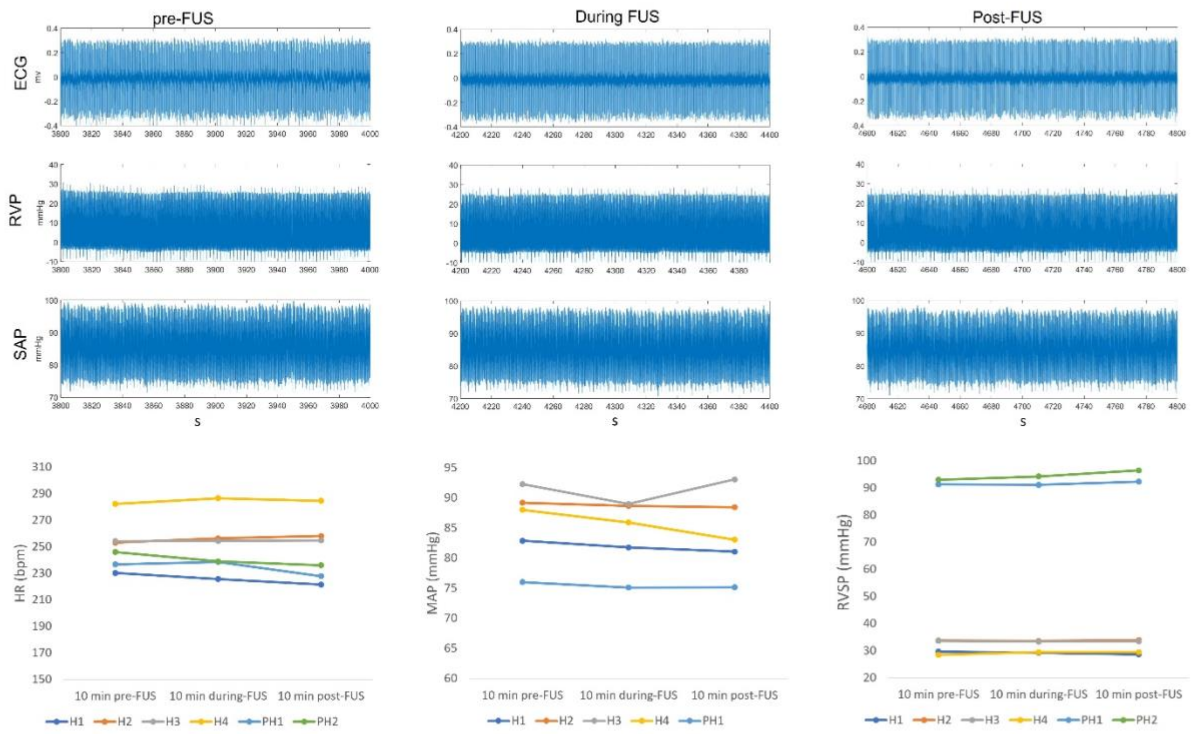


**Suppl. Fig. S3.** Representative examples of real-time right ventricular systolic pressure, arterial pressure, and heart rate raw recordings pre-, during- and post-FUS. The acute hemodynamic response to focused ultrasound of the spleen was recorded in 4 healthy animals (H1, H2, H3, H4) and 2 animals with pulmonary hypertension (PH1, PH2). HR, MAP and RVSP were averaged over a period of 10 min pre-FUS, during-FUS, and post-FUS.

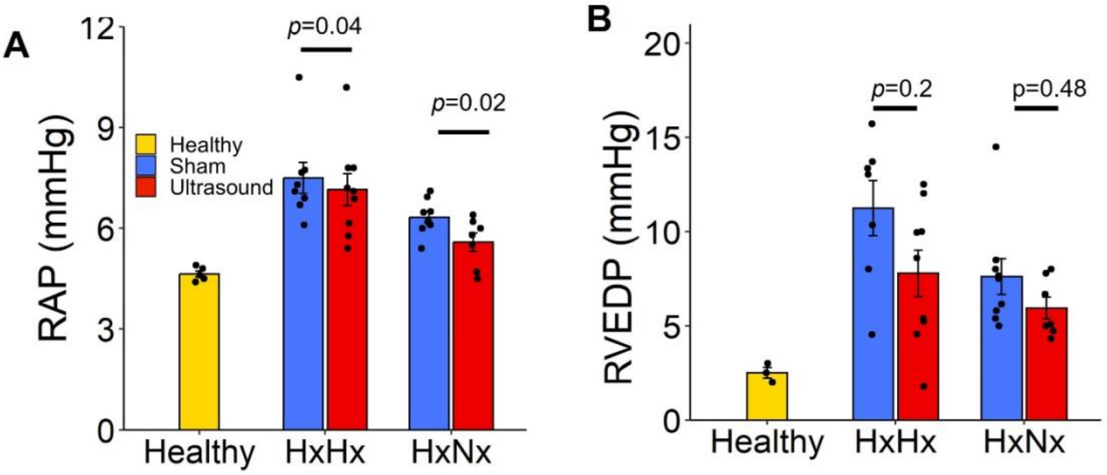


**Suppl. Fig. S4. (A)** Mean (±SEM) right atrial pressure (RAP) in HxHx and HxNx cohorts **(B)** Mean (±SEM) right ventricular end-diastolic pressure (RVEDP) in HxHx and HxNx cohorts.

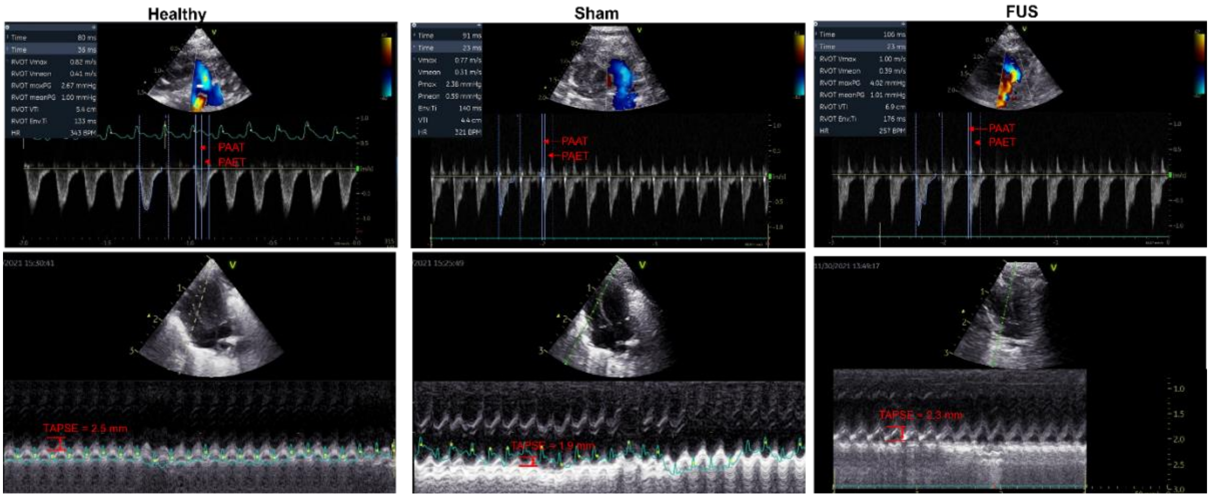


**Suppl. Fig. S5.** (A) Pulsed-wave doppler in the right ventricular outflow tract to measure pulmonary artery acceleration time (PAAT) and pulmonary artery ejection time (PAET) and (B) apical four-chamber ultrasound views, placing the M-mode line at the lateral tricuspid valve annulus to measure tricuspid annular plane systolic excursion in a healthy (left), sham-stimulated (middle) and sFUS (right) animals.

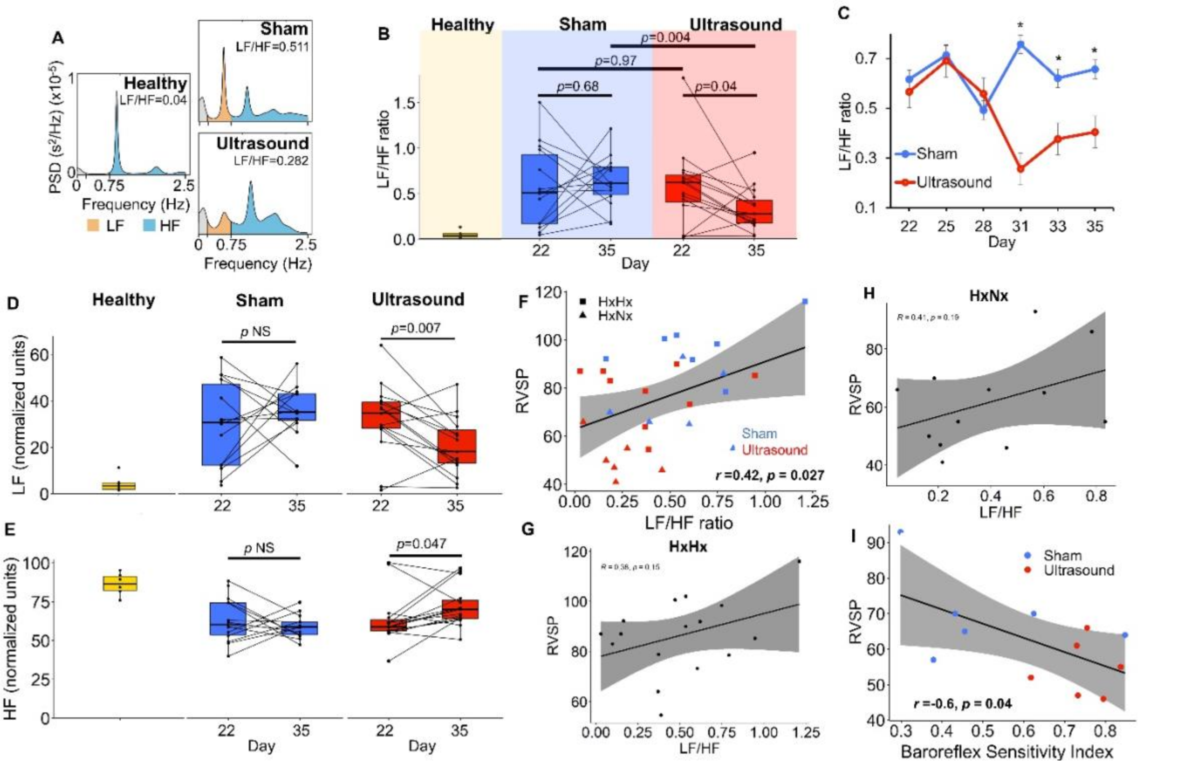


**Suppl. Fig. S6. sFUS improves indices of autonomic function in rats with PH.**
(A) Example heart rate power spectra used in the frequency domain analysis of heart rate variability (HRV) (13) in a healthy, sham-treated and sFUS-treated animal. The low-frequency (LF) component (0.2 to 0.75 Hz, shaded in orange) is considered an index of sympathetic activity, whereas the high-frequency (HF) component (0.75 to 2.5 Hz, shaded in blue) an index of parasympathetic activity. The LF/HF ratio is a metric of HRV: small LF/HF ratio is associated with a “healthier” sympathetic-parasympathetic balance. (B) LF/HF ratio on the first (day 22) and last day (day 35) of sFUS (n=15 animals) or sham treatment (n=14). LF/HF ratio is similar in the sFUS and sham groups on day 22 (p=0.97, unpaired t-test) and is reduced in sFUS on day 35 (p=0.004, unpaired t-test). LF/HF ratio does not change between day 22 and day 35 in the sham group (p=0.68, paired t-test), while it decreases in the sFUS group (p=0.04, paired t-test). (C) Time course of LF/HF ratio (mean±SE) throughout the treatment period, in sFUS- and sham-treated animals. Asterisks at different time points represent significant difference in LF/HF ratio between the 2 groups ($p<0.05$, repeated measures ANOVA). (D) Mean (±SD) Low frequency on day 22 and day 35 in sham- (n=14) and sFUS-stimulated(n=15) animals. (E) Mean (±SD) high frequency on day 22 and day 35 in sham- (n=14) and sFUS-stimulated(n=15) animals. (F) Linear correlation and 95% confidence intervals (shaded area) between RVSP and LF/HF ratio in sFUS- (blue symbols) and sham-treated animals (red symbols), in the hypoxia-normoxia (HxNx, triangles) and hypoxia-hypoxia cohorts (HxHx, squares). Pearson correlation coefficient r=-0.42 (p=0.027). (G and H) Correlations of RVSP and LF/HF in HxHx and HxNx cohorts. (I) Linear correlation and 95% confidence intervals (shaded area) between RVSP and Baroreflex Sensitivity Index (Pearson correlation coefficient r=-0.6, p=0.04).

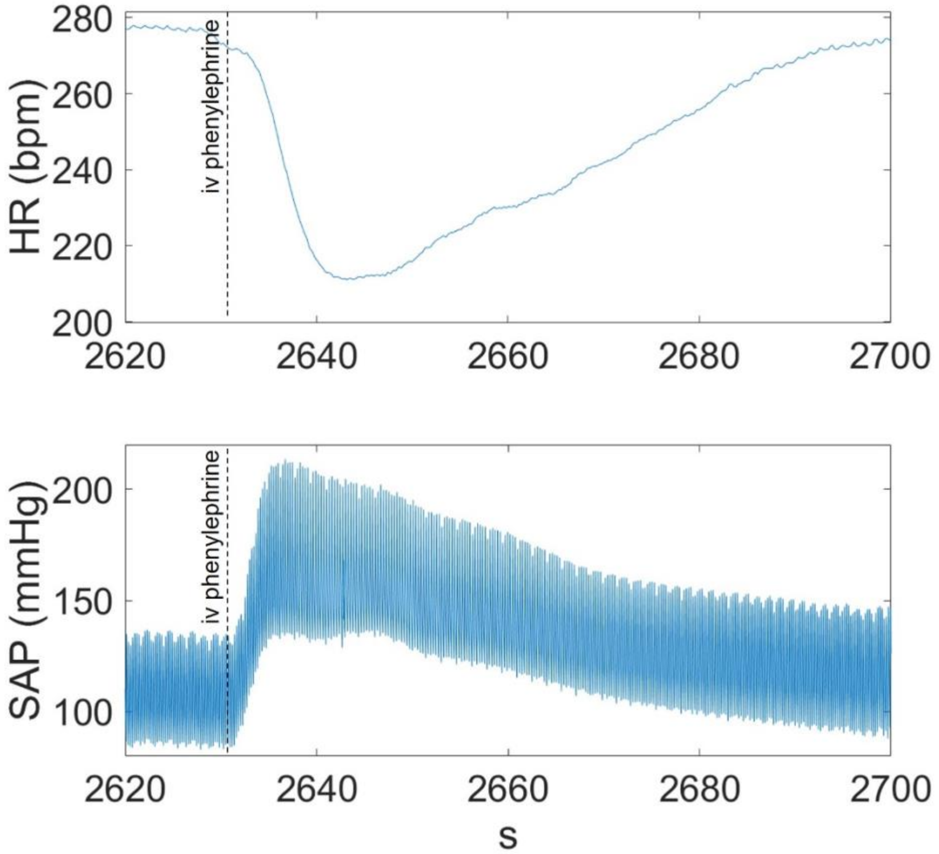


**Suppl. Fig. S7.** Representative example of baroreflex sensitivity index calculation in response to phenylephrine injection.

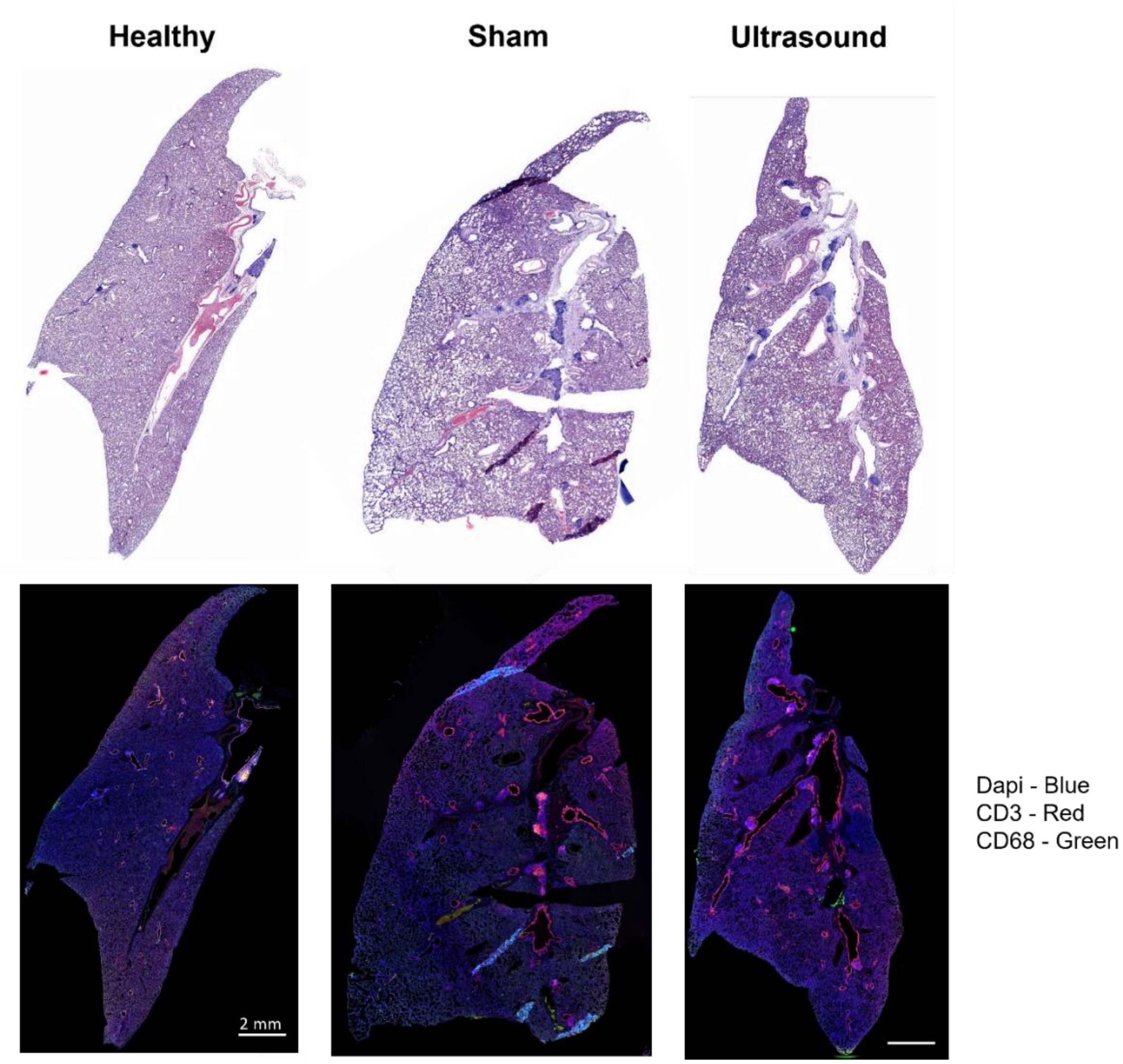


**Suppl. Fig. S8.** H&E staining and Immunohistochemistry in a healthy, sham-stimulated and sFUS animal.

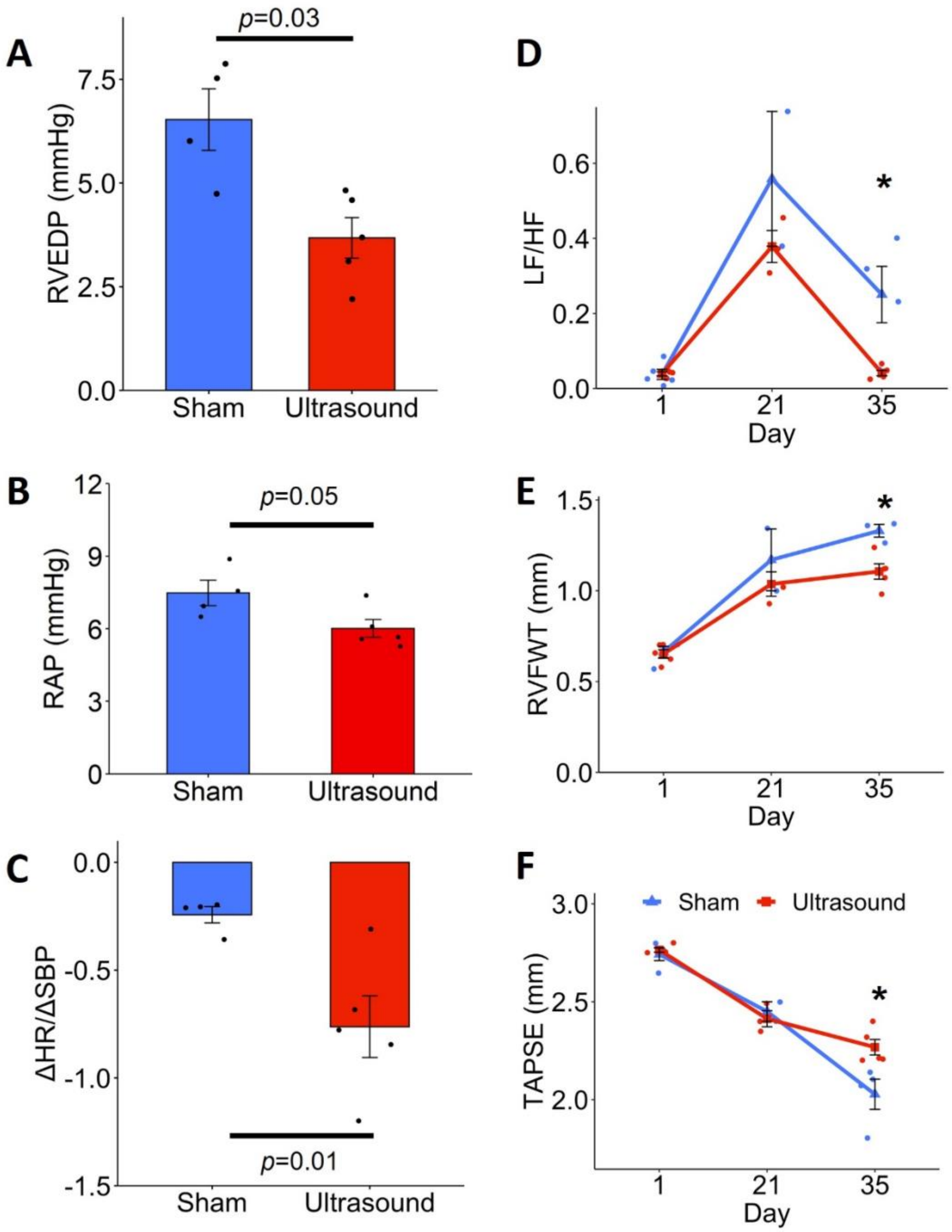


**Suppl. Fig. S9. Early and longer treatment cohort endpoints**

**(A)** Mean (±SEM) right ventricular end-diastolic pressure (RVEDP) in prevention cohort

**(B)** Mean (±SEM) right atrial pressure (RAP)

**(C)** Baroreflex sensitivity index calculated by change in heart rat over change in blood pressure after phenylephrine injection

**(D)** Line type plot of heart rate variability measured by low-frequency to high-frequency ratio (LF/HF) over time

**(E)** Line type plot of right ventricular free wall thickness (RVFWT) measured with echocardiography over time

**(F)** Line type plot of right ventricular function using tricuspid annular plane systolic excursion (TAPSE) measured in echocardiography over time

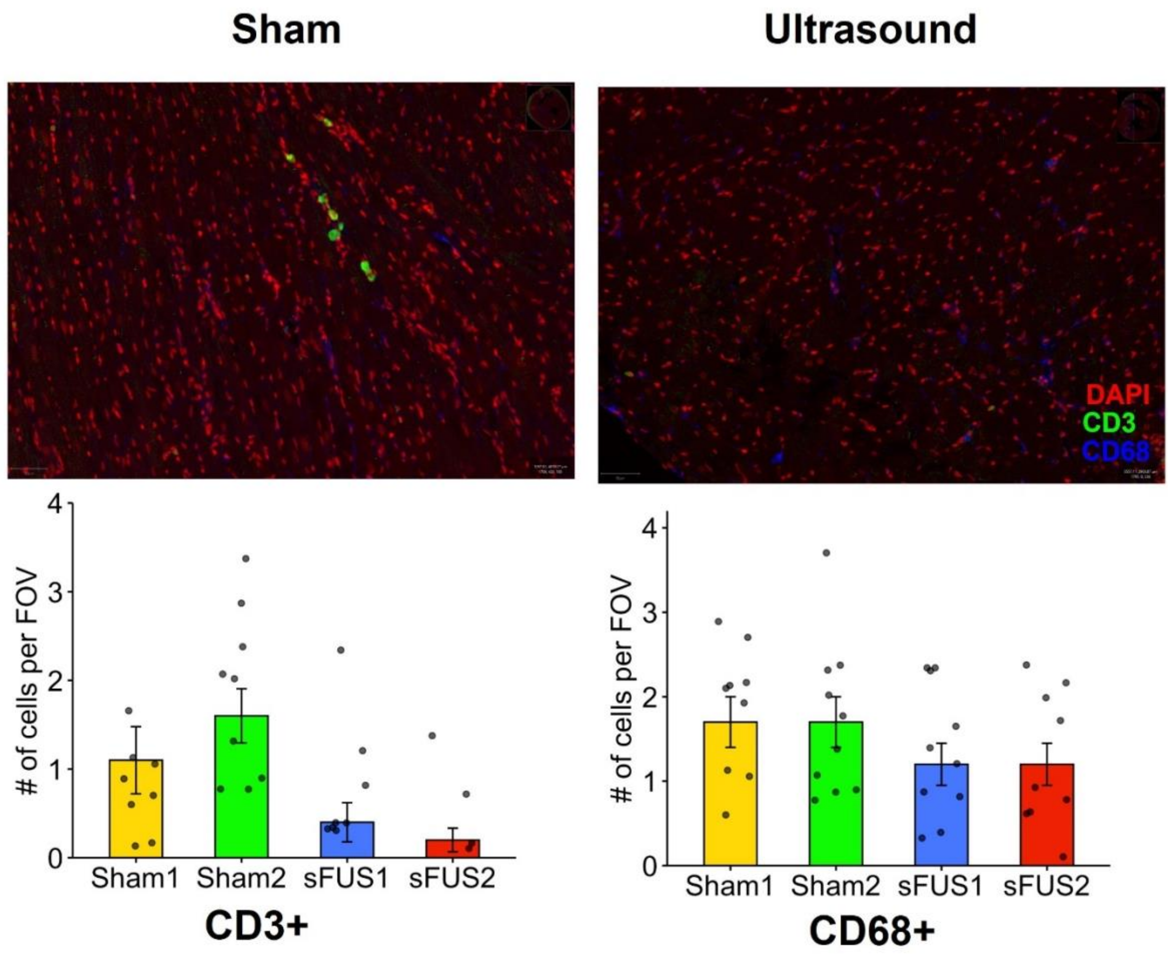


**Suppl. Fig. S10.** CD3+ and CD68+ cell counts in the hearts of 2 sham-stimulated and 2 sFUS animals.

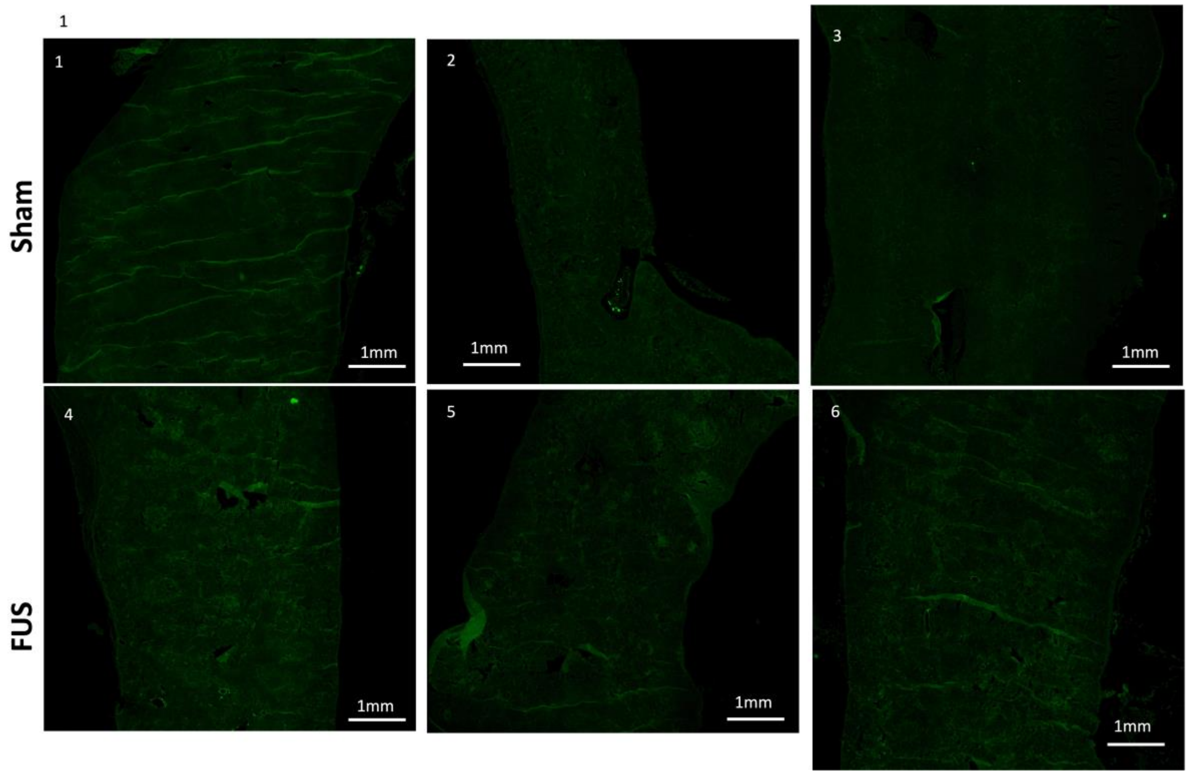


**Suppl. Fig. S11**. Spleen sections from 3 sham- and 3 ultrasound-stimulation rats with PH.

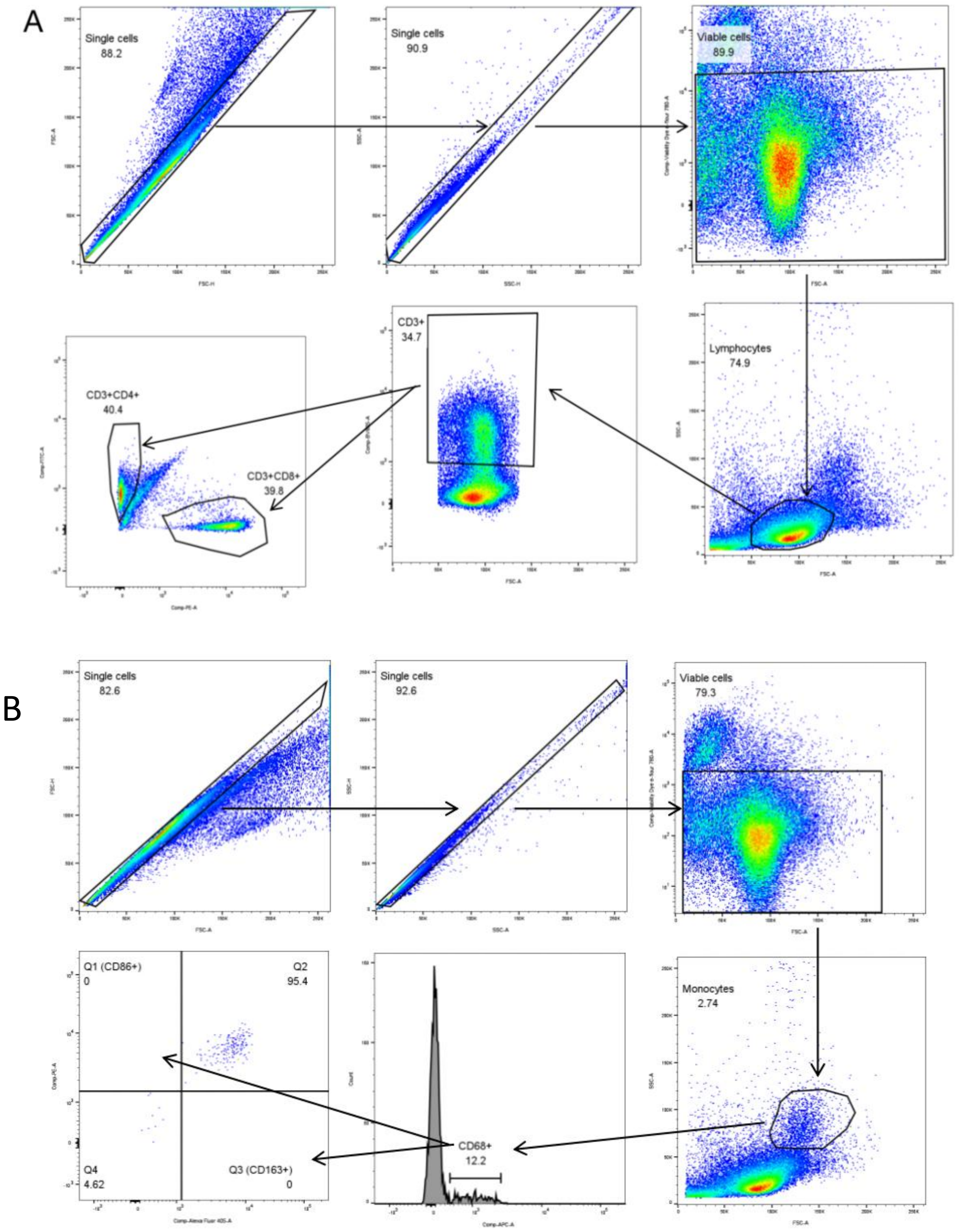


**Suppl. Fig. S12.** Gating strategies for (A) lymphocytes and (B) monocytes

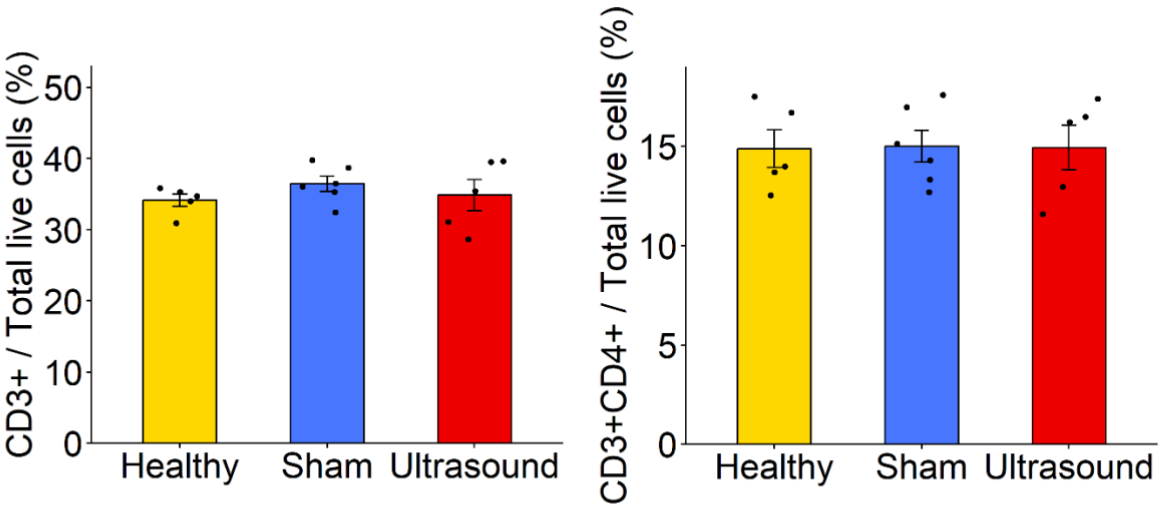


**Suppl. Fig. S13.** CD3+ and CD3+CD4+ cells in the spleen

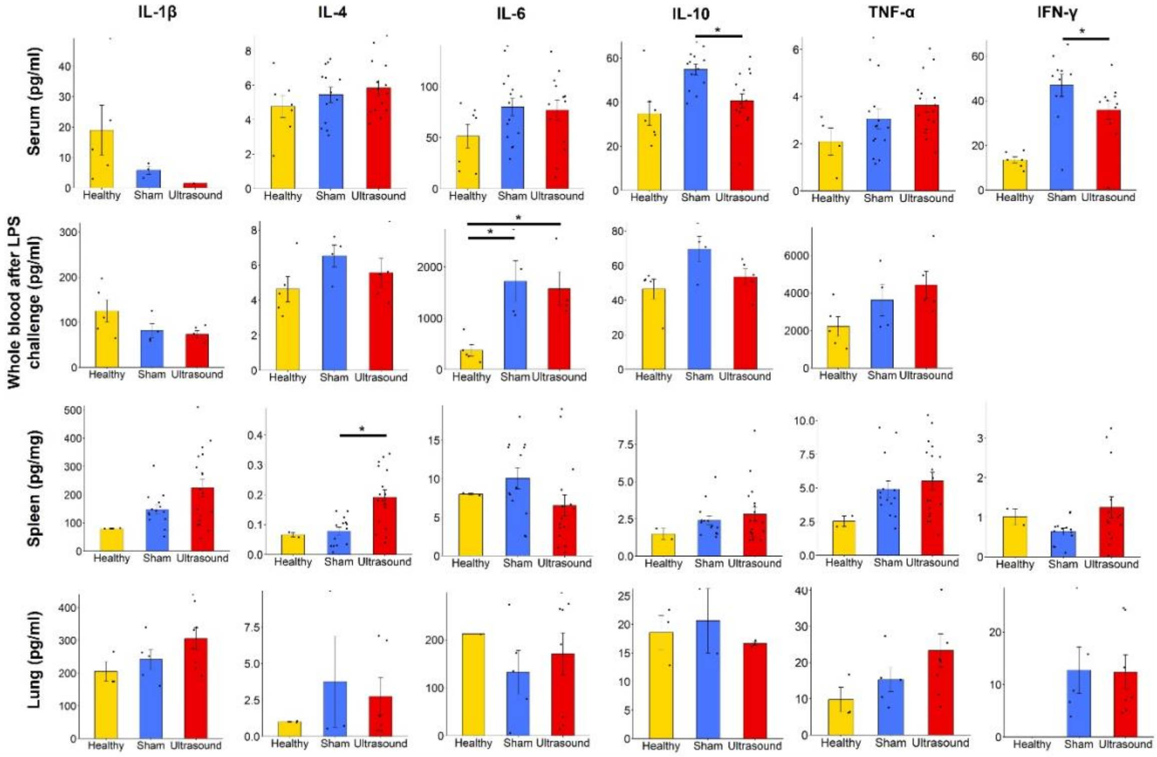


**Suppl. Fig. S14. Ultrasound effect on inflammatory cytokines.**
Mean (±SEM) of interleukin (IL)-1β, -4, -6, -10, tumor necrosis factor-a (TNF-a), and interferon-γ (IFN-γ) in healthy, sham-stimulated and ultrasound stimulated rats. The first row shows circulating cytokines values in the serum at day 35, the second row shows whole blood cytokines values after LPS ex-vivo challenge, the third row shows the cytokines values in spleen homogenates and the fourth row shows the cytokines values in spleen homogenates.

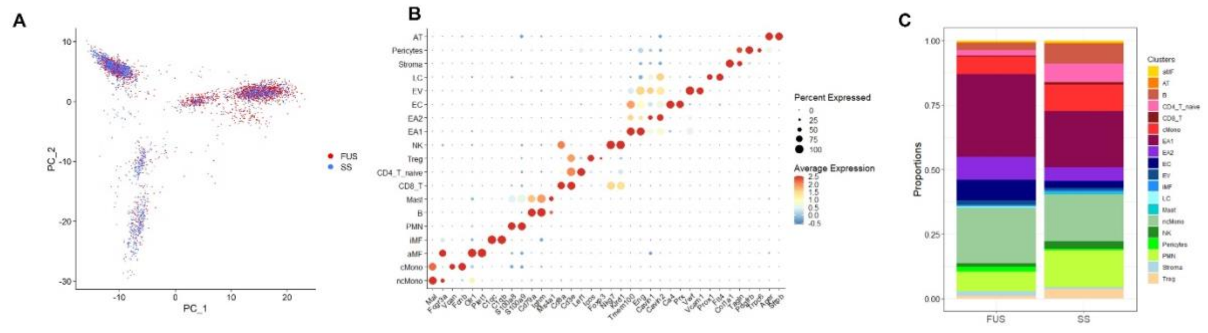

**Suppl. Fig. S15** (A) Principal component plot of lung cells from sham-stimulated (SS-blue) and focused ultrasound stimulated (FUS-red) rats. (B) Dot plot highlighting log10 average expression of 2 select marker genes used to identify cell clusters. The dot size corresponds to the percentage of cells expressing a gene in a given cluster. (C) Cell type proportions between FUS- and sham-stimulated rats.

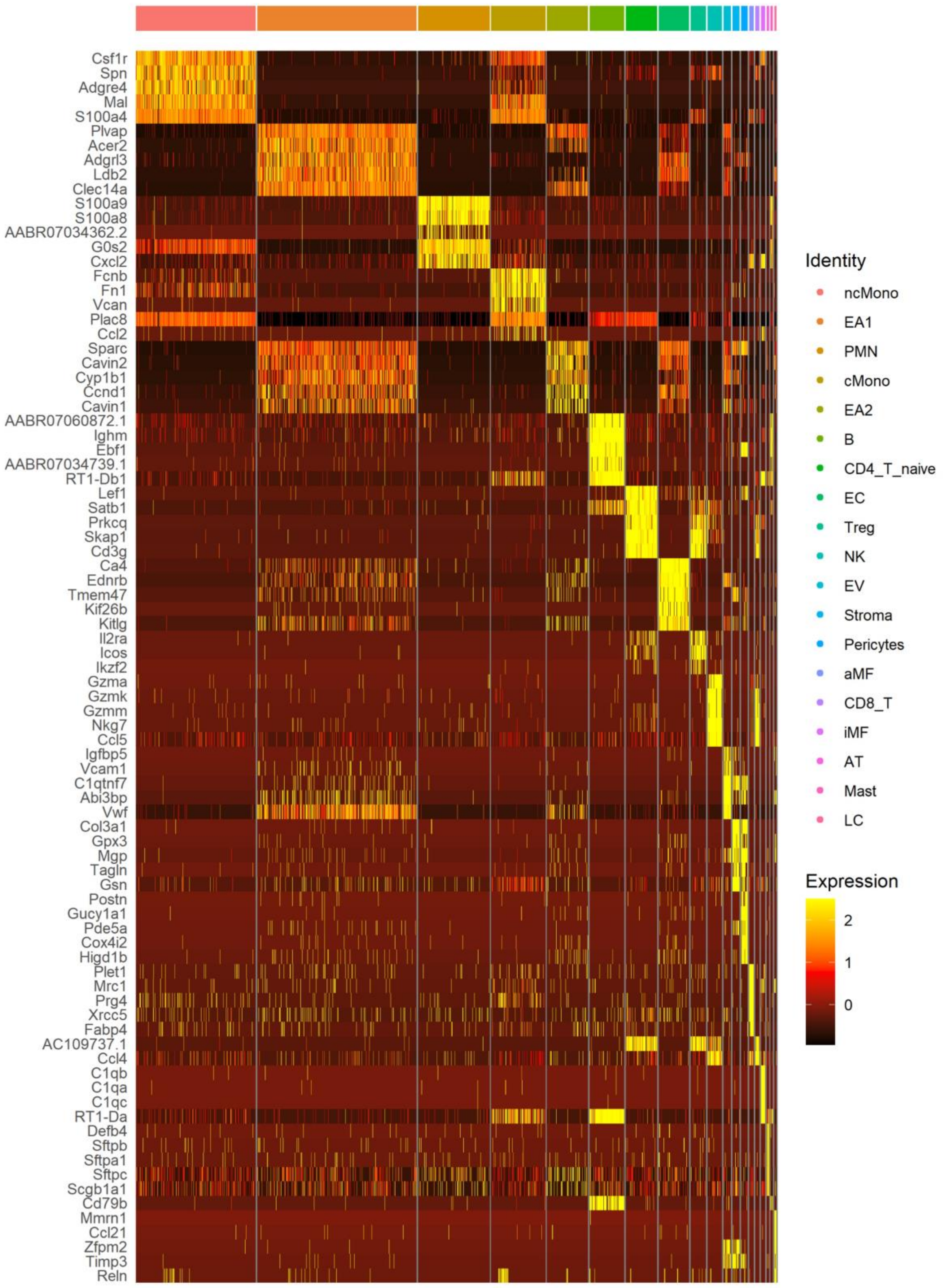


**Suppl. Fig. S16.** Heat map of RNA expression of the most selectively-expressed genes across all the cell types.

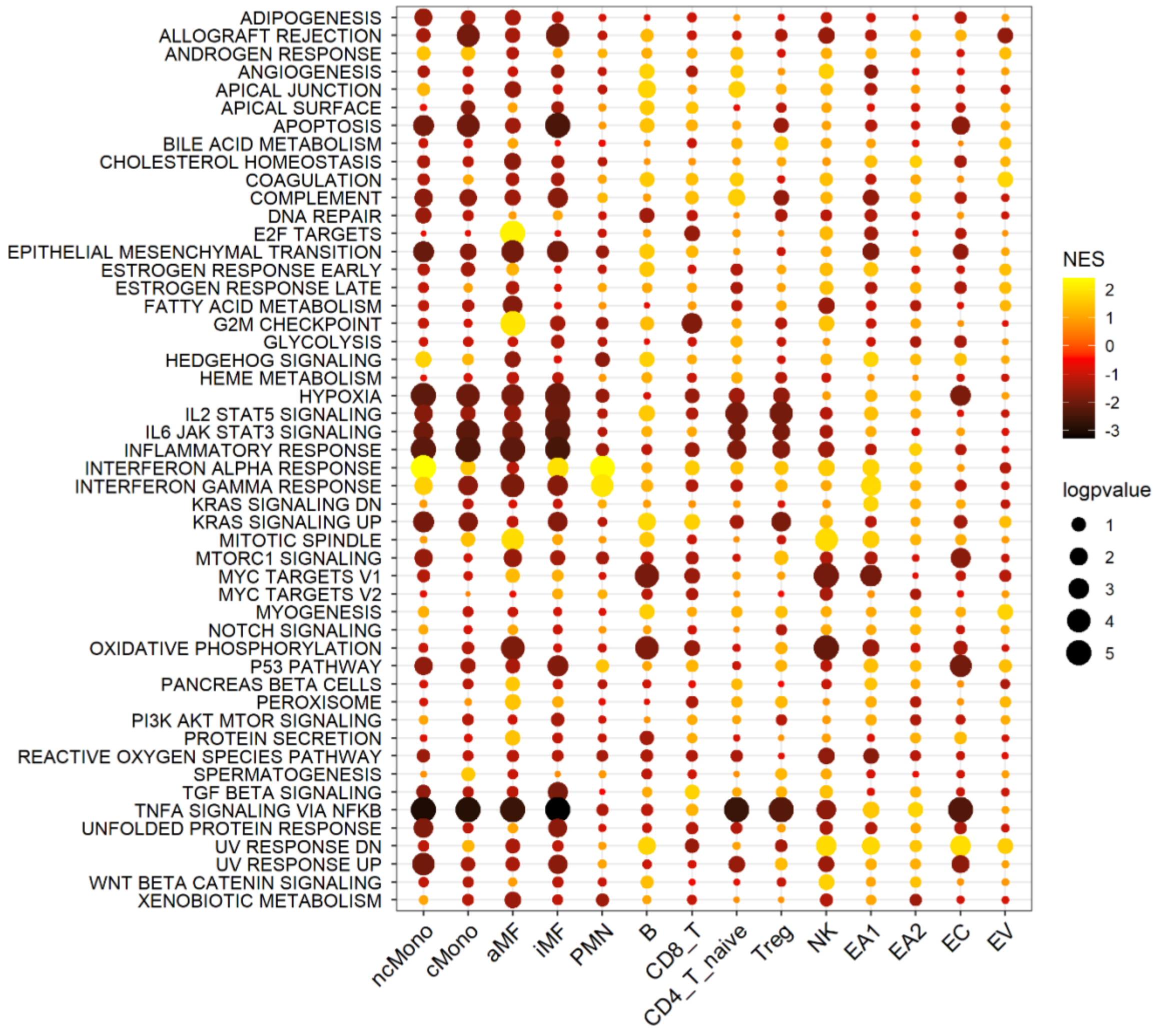


**Suppl. Fig. S17.** Heatmap showing all cell type-specific pathway enrichment between FUS and sham-stimulated rats using gene-set enrichment analysis (GSEA) (P<0.05) and hallmark pathways from the Molecular Signatures Database on the y-axis. The dot size corresponds to -log10(P), and color represents the normalized enrichment score (NES) from GSEA, indicating upregulation (yellow) or downregulation (black). TNFa/NF-kB signaling was significantly downregulated across many cell types while interferon alpha response was the most upregulated pathway.

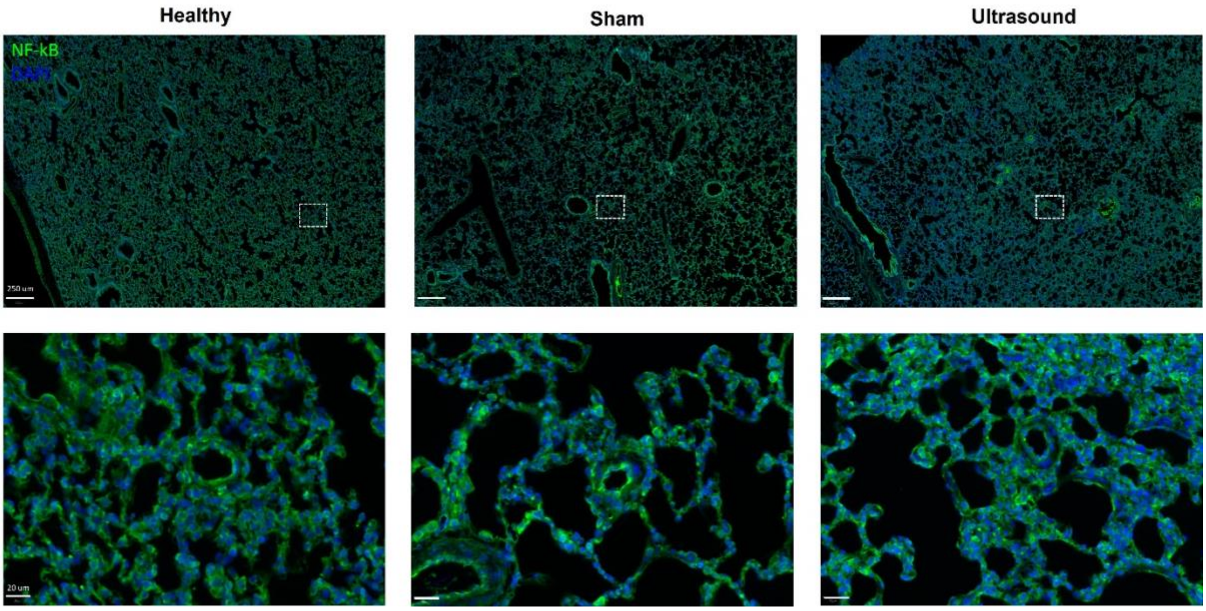


**Suppl. Fig. S18.** Representative images of nuclear staining NF-κB p65 and DAPI in lungs from a healthy, sham- and sFUS-treated rat.

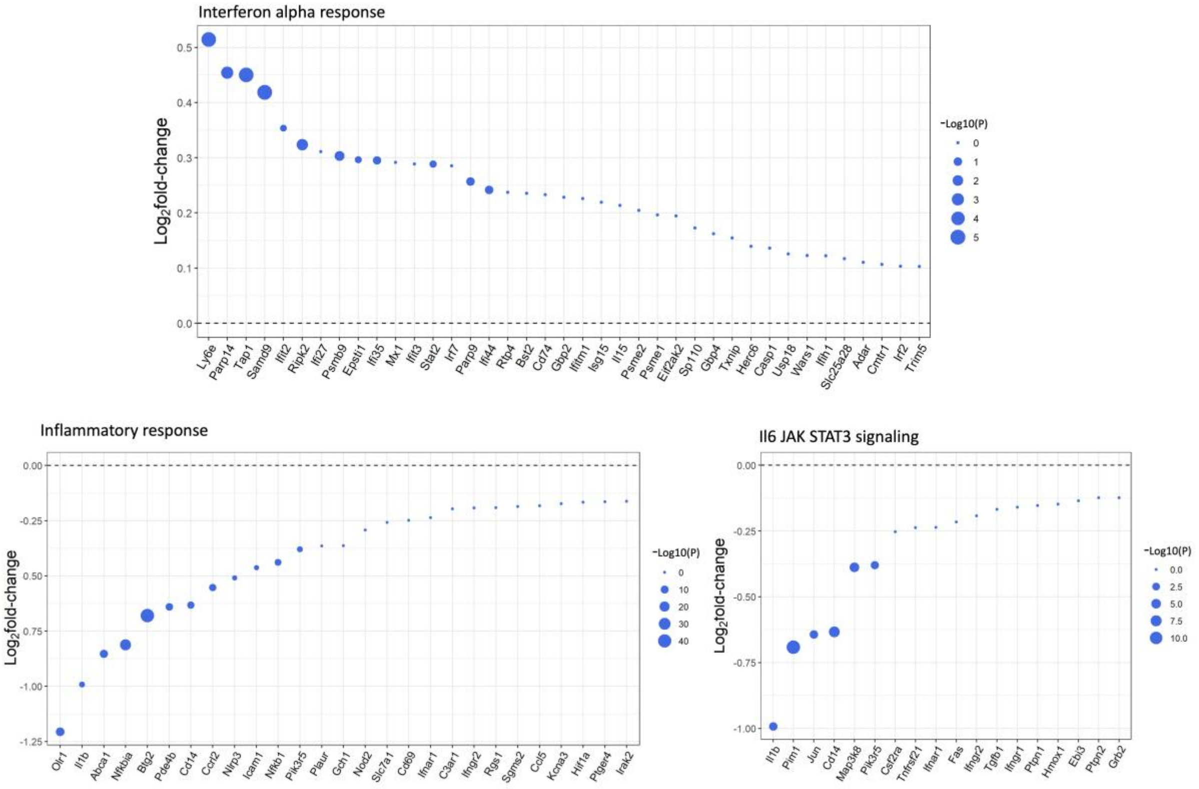


**Suppl. Fig. S19.** Dot plots showing MAST (Model-based Analysis of Single-Cell Transcriptomics) log2fold change of leading-edge genes accounting for the nonclassical monocytes upregulation of the IFNa response and downregulation of the inflammatory response and IL-6 JAK STAT3 signaling pathway as determined by GSEA, in which the size and color tint of dots represent the -log10(P-value).

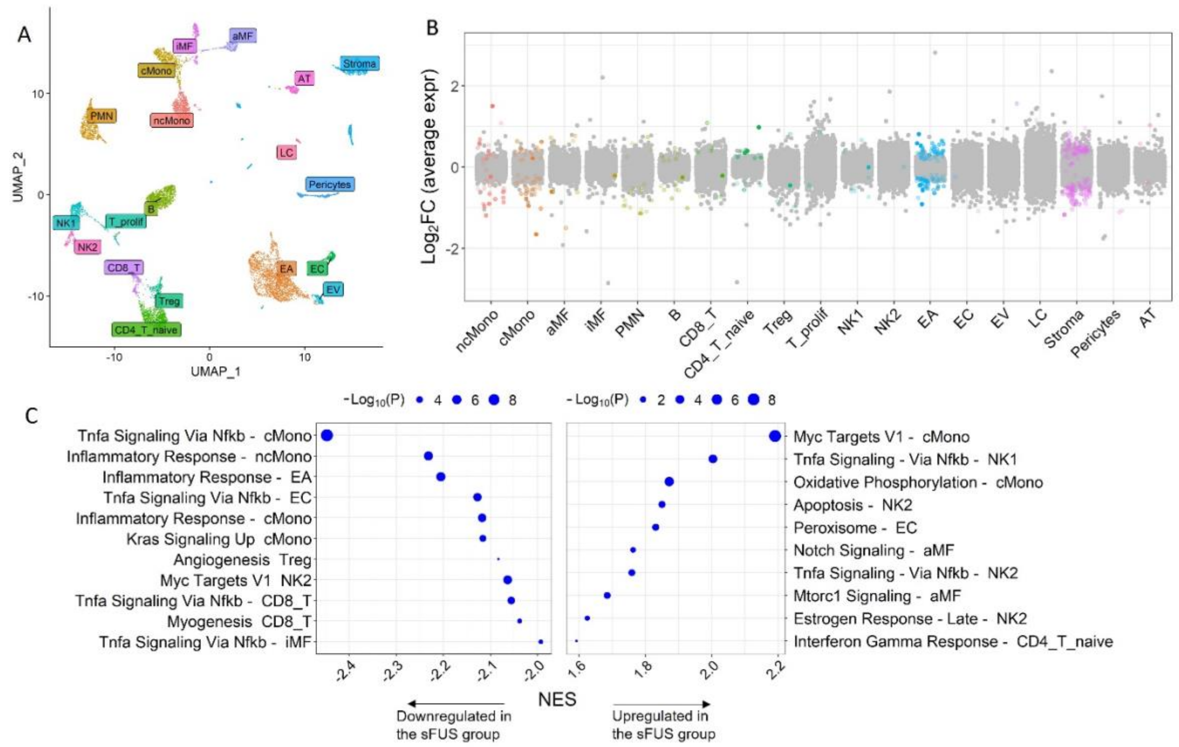


**Suppl. Fig. S20. ScRNA-seq of the lungs from rats with PH that received sFUS or sham stimulation in the "early and longer treatment cohort" (n=2/group). (A)** Uniform manifold approximation and projection (UMAP) plot showing lung cells from 4 rats with clusters labeled by cell type. **(B)** Jitter plot showing changes in gene expression for each cell type between sFUS and sham-stimulated rats. Each dot represents the differential expression MAST (Model-based Analysis of Single-Cell Transcriptomics) Log2-fold change of a gene. Dots indicating an adjusted p-value<0.05 are in color. The gray dots indicate values that were not significant (ns). cell type-specific pathway enrichment between sFUS and sham-stimulated rats using gene-set enrichment analysis (GSEA) (P<0.05) and 10 hallmark pathways from the Molecular Signatures Database that are known to be implicated in PAH on the y-axis. The dot size corresponds to -log10(P), and color represents the normalized enrichment score (NES) from GSEA, indicating upregulation (yellow) or downregulation (black). TNFa/NF-kB signaling was significantly downregulated across many cell types while interferon alpha response was the most upregulated pathway. **(C)** Dot plot showing the 10 most cell-specific downregulated (left) and upregulated pathways (right), using gene-set enrichment analysis (GSEA) (P<0.05) and the hallmark pathways from the Molecular Signatures Database that are on the y-axis. The dot size corresponds to -log10(P) and x-axis represents the normalized enrichment score (NES) from GSEA.

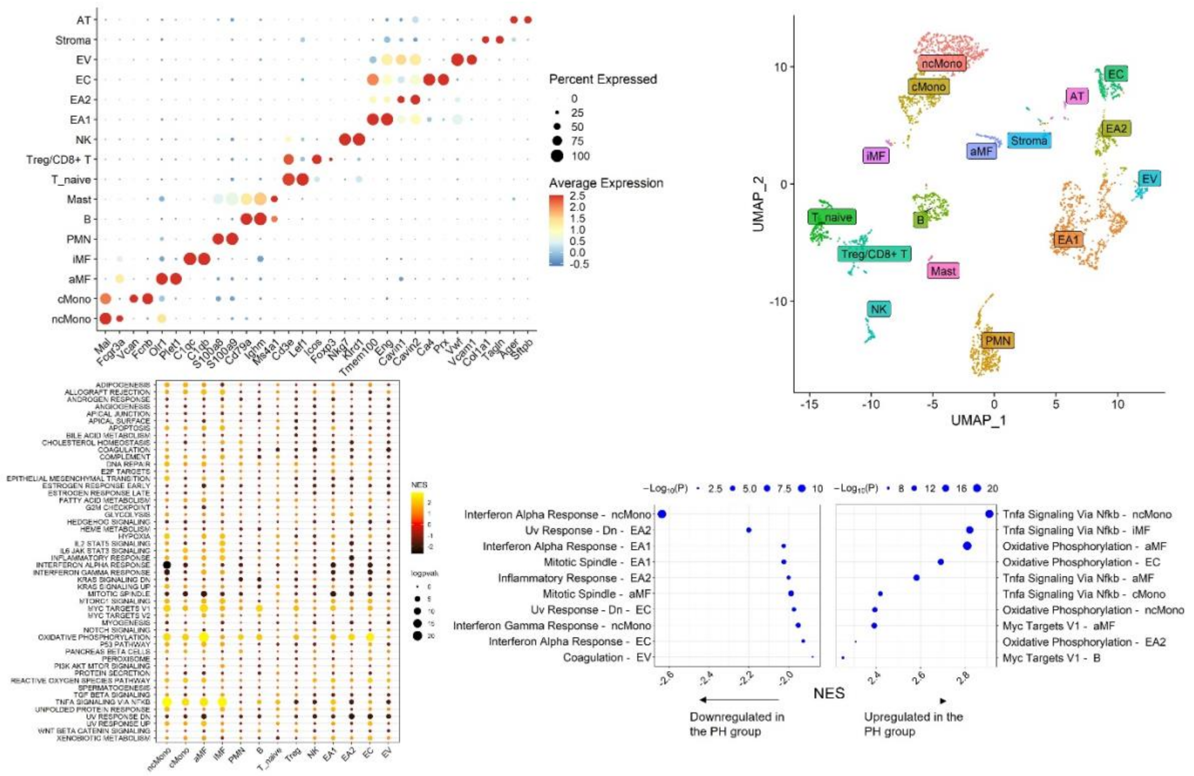


**Suppl. Fig. S21**

**ScRNA-seq of the lungs from healthy rats and rats with PH (n=2/group).** (A) Dot plot highlighting log10 average expression of 2 select marker genes used to identify cell clusters. The dot size corresponds to the percentage of cells expressing a gene in a given cluster. (B) Uniform manifold approximation and projection (UMAP) plot showing lung cells from 4 rats with clusters labeled by cell type. (C) Heatmap showing major cell type-specific pathway enrichment between FUS and sham-stimulated rats using gene-set enrichment analysis (GSEA) (P<0.05) and 10 hallmark pathways from the Molecular Signatures Database that are known to be implicated in PAH on the y-axis. The dot size corresponds to -log10(P), and color represents the normalized enrichment score (NES) from GSEA, indicating upregulation (yellow) or downregulation (black). (D) Dot plot showing the 10 most cell-specific downregulated (left) and upregulated pathways (right).

### Supplementary tables

| Cohort | Characteristic | Number of animals | Readouts | Fig. |
|---|---|---|---|---|
| Pre-emptive splenic denervation | Splenic denervation (or sham surgery) before SuHxNx induction (or normoxia) | PH-/SDN-: 5<br>PH-/SDN+: 5<br><br>PH+/SDN-: 4<br><br>PH+/SDN-: 5 | Hemodynamics, echocardiography, lung histology | 1 |
| **SUGEN-Hypoxia model** | | | | |
| Treatment<br>• HxHx | Ultrasound or sham stimulation | Sham: 8<br><br>sFUS: 9 | Hemodynamics, autonomics, cytokines, BNP, lung histology | 2, 3 |
| • HxNx | Ultrasound or sham stimulation | Sham: 8<br><br>sFUS: 8 | Hemodynamics, autonomics, cytokines, BNP, echocardiography, lung histology and IHC, flow cytometry, scRNA-seq | 2, 3 |
| Extended follow-up | Ultrasound or sham stimulation | Sham: 5<br><br>sFUS: 7 | Hemodynamics, autonomics, echocardiography, lung histology | 5 |
| Early treatment | Ultrasound or sham stimulation | Sham: 4<br><br>sFUS: 5 | Hemodynamics, autonomics, cytokines, BNP, echocardiography, lung and heart histology and IHC, scRNA-seq | 5 |
| Splenic denervation | Ultrasound or sham stimulation | Sham: 4<br><br>sFUS: 4 | Hemodynamics, autonomics, echocardiography, lung histology | 6 |
| **MCT model** | | | | |
| | Ultrasound or sham stimulation | Sham: 7<br><br>sFUS: 7 | Hemodynamics, autonomics, echocardiography, lung histology | 4 |

**Supplemental Table S1. Summary of all cohorts.**

| Characteristic | **Sham**, N = 5 | **Ultrasound**, N = 6 | p-value |
|---|---|---|---|
| **Max dp/dt, mmHg/s** | 2,754 (221) | 2,236 (217) | **0.042** |
| **Min dP/dt, mmHg/s** | -2,780 (340) | -2,035 (304) | **0.024** |
| **Diastolic duration, s** | 0.105 (0.008) | 0.108 (0.009) | 0.4 |
| **Systolic duration, s** | 0.098 (0.017) | 0.097 (0.011) | 0.7 |
| **Cycle duration, s** | 0.203 (0.023) | 0.205 (0.019) | 0.8 |
| **Contractility index, 1/s** | 111 (12) | 114 (21) | >0.9 |
| **Tau, s** | 0.027 (0.011) | 0.040 (0.015) | 0.3 |
| **Pressure Time index, mmHg.s** | 4.49 (1.34) | 3.44 (0.72) | 0.2 |
| **PAAT, ms** | 16.67 (0.80) | 18.33 (0.67) | 0.14 |
| **PAAT/PAET** | 0.19 (0.01) | 0.21 (0.01) | 0.28 |
| **TAPSE, mm** | 2.37 (0.12) | 2.56 (0.12) | 0.3 |
| Mean(SD) | | | |

**Supplemental Table S2. RVSP waveform analysis and echocardiographic parameters in HxNx cohort.**

| Characteristic | **Sham** (n = 14) | **Ultrasound** (n = 15) | p-value |
|---|---|---|---|
| **SDSD** | 1.73(0.72) | 2.33(1.36) | 0.2 |
| **RMSSD** | 1.73(0.72) | 2.33(1.36) | 0.2 |
| **LF Power, ms$^2$** | 0.68(0.54) | 0.65(0.62) | 0.7 |
| **LF Power, %** | 26(12) | 16(11) | **0.037** |
| **LF Power, nu** | 32(12) | 19(12) | **0.006** |

| | | | |
|---|---|---|---|
| **HF Power, ms²** | 0.98(0.59) | 2.54(2.65) | 0.057 |
| **HF Power, %** | 42(13) | 57(13) | **0.003** |
| **HF Power, nu** | 57(7) | 68(13) | **0.010** |
| Mean(SD) | | | |

**Supplemental Table S3. Time- and frequency-domain heart rate variability metrics.**

| Characteristic | **Sham**, N = 5 | **Ultrasound**, N = 5 | p-value |
|---|---|---|---|
| **PAAT, ms** | 17.67 (0.67) | 20.1 (0.81) | 0.08 |
| **PAAT/PAET** | 0.20 (0.01) | 0.23 (0.01) | 0.1 |
| **TAPSE, mm** | 2.47 (0.8) | 2.66 (0.10) | 0.19 |
| Mean(SE) | | | |

**Supplemental Table S4. Echocardiographic parameters in MCT model.**

| Characteristic | **Sham**, N = 4 | **Ultrasound**, N = 5 | p-value |
|---|---|---|---|
| **Max dp/dt, mmHg/s** | 3,890 (766) | 2,789 (545) | **0.037** |
| **Min dP/dt, mmHg/s** | -3,938 (778) | -2,692 (774) | **0.033** |
| **Diastolic duration, s** | 0.107 (0.012) | 0.116 (0.016) | 0.4 |
| **Systolic duration, s** | 0.090 (0.005) | 0.096 (0.008) | 0.4 |
| **Cycle duration, s** | 0.197 (0.015) | 0.212 (0.022) | 0.4 |
| **Contractility index, 1/s** | 179 (81) | 141 (30) | 0.7 |
| **Tau, s** | 0.039 (0.021) | 0.037 (0.010) | 0.7 |
| **Pressure Time index, mmHg.s** | 5.52 (0.54) | 4.37 (0.70) | 0.016 |
| Mean(SD) | | | |

**Supplemental Table S5. RVSP waveform analysis in early treatment cohort.**

| Characteristic | **Healthy**, N = 7 | **Sham**, N = 15 | **Ultrasound**, N = 17 | p-value[1] |
|---|---|---|---|---|
| **IFN-γ** | 14 (1) | 47 (5) | 36 (4) | 0.003 |
| **IL-10** | 35 (5) | 55 (2) | 41 (3) | 0.002 |
| **IL-13** | 12.5 (1.5) | 8.6 (1.3) | 7.2 (1.1) | 0.054 |
| **IL-1β** | 19 (8) | 6 (1) | 1 (NA) | 0.2 |
| **IL-4** | 4.77 (0.64) | 5.44 (0.44) | 5.84 (0.43) | 0.5 |
| **IL-5** | 12.4 (3.1) | 15.3 (3.0) | 15.5 (1.7) | 0.5 |
| **IL-6** | 51 (12) | 80 (9) | 77 (10) | 0.2 |
| **KC/GRO** | 153 (21) | 164 (25) | 176 (39) | 0.9 |
| **TNF-α** | 2.09 (0.58) | 3.06 (0.41) | 3.64 (0.31) | 0.090 |

[1] Kruskal-Wallis rank sum test

**Supplemental Table S6. Inflammatory cytokines measured in serum from healthy (n=7), sham- (n=15) and sFUS (n=17) rats**

| Characteristic | **Healthy**, N = 5 | **Sham**, N = 4 | **Ultrasound**, N = 5 | p-value[1] |
|---|---|---|---|---|
| **IL-10** | 46 (6) | 70 (7) | 53 (5) | 0.2 |
| **IL-13** | 10.2 (3.2) | 11 (2) | 4 (2) | 0.2 |
| **IL-1β** | 125 (25) | 82 (15) | 73 (8) | 0.2 |
| **IL-4** | 4.63 (0.72) | 6.53 (0.62) | 5.57 (0.81) | 0.2 |
| **IL-5** | 15 (3) | 13 (8) | 21 (4) | 0.6 |
| **IL-6** | 373 (110) | 1,720 (394) | 1,574 (330) | 0.008 |
| **KC/GRO** | 108 (28) | 215 (32) | 775 (647) | 0.12 |

| | | | | |
|---|---|---|---|---|
| **TNF-α** | 2,210 (523) | 3,619 (847) | 4,426 (721) | 0.1 |
| [1] Kruskal-Wallis rank sum test | | | | |

**Supplemental Table S7. Inflammatory cytokines measured in whole blood after ex-vivo LPS challenge from healthy (n=6), sham- (n=4) and sFUS (n=5) rats**

| **Characteristic** | **Healthy**, N = 2 | **Sham**, N = 14 | **Ultrasound**, N = 18 | **p-value**[1] |
|---|---|---|---|---|
| **IFN-γ** | 1.01 (0.20) | 0.64 (0.08) | 1.25 (0.27) | 0.09 |
| **IL-10** | 1.49 (0.36) | 2.38 (0.29) | 2.81 (0.45) | 0.5 |
| **IL-13** | 0.20 (0.00) | 0.41 (0.08) | 0.52 (0.14) | 0.7 |
| **IL-1β** | 80 (1) | 148 (16) | 225 (31) | 0.10 |
| **IL-4** | 0.07 (0.01) | 0.08 (0.01) | 0.19 (0.02) | 0.005 |
| **IL-5** | 1.83 (0.10) | 2.68 (0.34) | 3.82 (0.46) | 0.09 |
| **IL-6** | 8.0 (0.1) | 10.1 (1.4) | 6.5 (1.4) | 0.13 |
| **KC/GRO** | 12 (4) | 38 (6) | 54 (10) | 0.10 |
| **TNF-α** | 2.54 (0.39) | 4.91 (0.60) | 5.51 (0.65) | 0.2 |
| [1] Kruskal-Wallis rank sum test | | | | |

**Supplemental Table S8. Inflammatory cytokines measured in spleen homogenates from healthy (n=2), sham- (n=14) and sFUS (n=18) rats**

| **Characteristic** | **Healthy**, N = 3 | **Sham**, N = 5 | **Ultrasound**, N = 8 | **p-value**[1] |
|---|---|---|---|---|
| **IFN-γ** | NA | 13 (4) | 12 (3) | >0.9 |
| **IL-10** | 19 (3) | 20.6 (5.6) | 17 (0) | 0.9 |
| **IL-13** | 2.25 (1.76) | 2.48 (0.80) | 3.60 (0.91) | 0.4 |
| **IL-1β** | 205 (30) | 242 (31) | 305 (33) | 0.2 |
| **IL-4** | 1.01 (0.04) | 3.76 (3.12) | 2.75 (1.28) | 0.9 |

| | | | | |
|---|---|---|---|---|
| **IL-5** | NA | 28 (4) | 24 (9) | 0.3 |
| **IL-6** | 212 (NA) | 133 (45) | 171 (44) | 0.7 |
| **KC/GRO** | 355 (78) | 707 (129) | 718 (97) | 0.083 |
| **TNF-α** | 10 (3) | 15 (3) | 23 (5) | 0.11 |

[1] Kruskal-Wallis rank sum test

**Supplemental Table S9. Inflammatory cytokines measured in lung homogenates from healthy (n=3), sham- (n=5) and sFUS (n=8) rats**
